# High-altitude adaptation and incipient speciation in geladas

**DOI:** 10.1101/2021.09.01.458582

**Authors:** Kenneth L. Chiou, Mareike C. Janiak, India Schneider-Crease, Sharmi Sen, Ferehiwot Ayele, Idrissa S. Chuma, Sascha Knauf, Alemayehu Lemma, Anthony V. Signore, Anthony M. D’Ippolito, Belayneh Abebe, Abebaw Azanaw Haile, Fanuel Kebede, Peter J. Fashing, Nga Nguyen, Colleen McCann, Marlys L. Houck, Jeffrey D. Wall, Andrew S. Burrell, Christina M. Bergey, Jeffrey Rogers, Jane E. Phillips-Conroy, Clifford J. Jolly, Amanda D. Melin, Jay F. Storz, Amy Lu, Jacinta C. Beehner, Thore J. Bergman, Noah Snyder-Mackler

**Affiliations:** Center for Evolution and Medicine, Arizona State University, Tempe, AZ, USA; School of Life Sciences, Arizona State University, Tempe, AZ, USA; Department of Psychology, University of Washington, Seattle, WA, USA; Nathan Shock Center of Excellence in the Basic Biology of Aging, University of Washington, Seattle, WA, USA; School of Science, Engineering, & Environment, University of Salford, Salford, UK; Department of Anthropology, University of Michigan, Ann Arbor, MI, USA; College of Veterinary Medicine and Agriculture, Addis Ababa University, Debre Zeit, Ethiopia; Veterinary Unit, Conservation Science Department, Tanzania National Parks (TANAPA), Arusha, Tanzania; Work Group Neglected Tropical Diseases, Infection Biology Unit, German Primate Center, Leibniz Institute for Primate Research, Göttingen, Germany; Institute for International Animal Health / One Health, Friedrich-Loeffler-Institut, Federal Research Institute for Animal Health, Greifswald, Island Riems, Germany; School of Biological Sciences, University of Nebraska, Lincoln, NE, USA; University Program in Genetics and Genomics, Duke University, Durham, NC, USA; Center for Genomic and Computational Biology, Duke University, Durham, NC, USA; African Wildlife Foundation, Debark, Ethiopia; Ethiopian Wildlife Conservation Authority, Addis Ababa, Ethiopia; Department of Anthropology and Environmental Studies Program, California State University Fullerton, Fullerton, CA, USA; Centre for Ecological and Evolutionary Synthesis, Department of Biosciences, University of Oslo, Oslo, Norway; Department of Mammals, Bronx Zoo, Wildlife Conservation Society, New York, NY, USA; New York Consortium in Evolutionary Primatology, New York, NY, USA; Beckman Center for Conservation Research, San Diego Zoo Wildlife Alliance, Escondido, CA, USA; Institute for Human Genetics, University of California San Francisco, San Francisco, CA, USA; Department of Anthropology, New York University, New York, NY, USA; Department of Genetics, Human Genetics Institute of New Jersey, Rutgers University, Piscataway, NJ, USA; Human Genome Sequencing Center, Baylor College of Medicine, Houston, TX, USA; Department of Neuroscience, Washington University School of Medicine, St. Louis, MO, USA; Department of Anthropology, Washington University, St. Louis, MO, USA; Department of Anthropology and Archaeology, University of Calgary, Calgary, AB, Canada; Alberta Children’s Hospital Research Institute, University of Calgary, Calgary, AB, Canada; Department of Medical Genetics, University of Calgary, Calgary, AB, Canada.; Department of Anthropology, Stony Brook University, Stony Brook, NY, USA; Department of Psychology, University of Michigan, Ann Arbor, MI, USA; Department of Ecology & Evolution, University of Michigan, Ann Arbor, MI, USA; Center for Studies in Demography & Ecology, University of Washington, Seattle, WA, USA

## Abstract

Survival at high altitude requires adapting to extreme conditions such as environmental hypoxia. To understand high-altitude adaptations in a primate, we assembled the genome of the gelada (*Theropithecus gelada*), an endemic Ethiopian monkey, and complemented it with population resequencing, hematological, and morphometric data. Unexpectedly, we identified a novel karyotype that may contribute to reproductive isolation between gelada populations. We also identified genomic elements including protein-coding sequences and gene families that exhibit accelerated changes in geladas and may contribute to high-altitude adaptation. Our findings lend insight into mechanisms of speciation and adaptation while providing promising avenues for functional hypoxia research.

---

Life at high altitude (≥2,500 meters) is associated with myriad environmental challenges including cold temperatures and reduced oxygen availability due to low barometric pressure. Consequently, organisms at high altitude have encountered strong evolutionary pressure to adapt to these challenges. Human populations living at high altitude, for example, have evolved physiological adaptations to hypoxia [1, 2], providing compelling examples of strong directional selection operating over short evolutionary time frames.

Human populations began living at high altitude quite recently, from as little as 150 years to as long as 47,000 years ago [3]. This time frame pales in comparison to that of nonhuman animals living at high altitude over macroevolutionary time (i.e., >1 million years). Such lineages would be expected to exhibit a greater number of fixed genetic and phenotypic differences relative to their closest lowland counterparts and provide a valuable comparative opportunity for understanding mechanisms underlying the evolution of high-altitude adaptations in humans and other animals. Comparative perspectives are particularly valuable for identifying both the shared and divergent routes that natural selection has taken at the nucleotide, protein, and pathway levels to facilitate adaptations to high-altitude life [4, 5].

The gelada (*Theropithecus gelada*) is a cercopithecoid monkey—closely related to baboons (*Papio* spp.) and *Lophocebus*/*Rungwecebus* mangabeys [7, 8]—endemic to Ethiopia (Fig. 1a–b). It is the only surviving member of the genus *Theropithecus*, which was found from South Africa to as far as Spain, Italy, and India up to 1 million years ago [9, 10]. Geladas likely avoided the fate of their extinct congenerics by exploiting an extreme environment over the past several million years: the grassy plateaus of the Ethiopian highlands [11]. Consequently, geladas have adopted primarily grass-eating diets and are found mainly at elevations from 2,350 to 4,550 meters above sea level (Fig. 1c) [12], representing one of the highest altitudinal ranges of any extant primate species, matched only by some *Rhinopithecus* monkeys [13].

**Figure 1.**
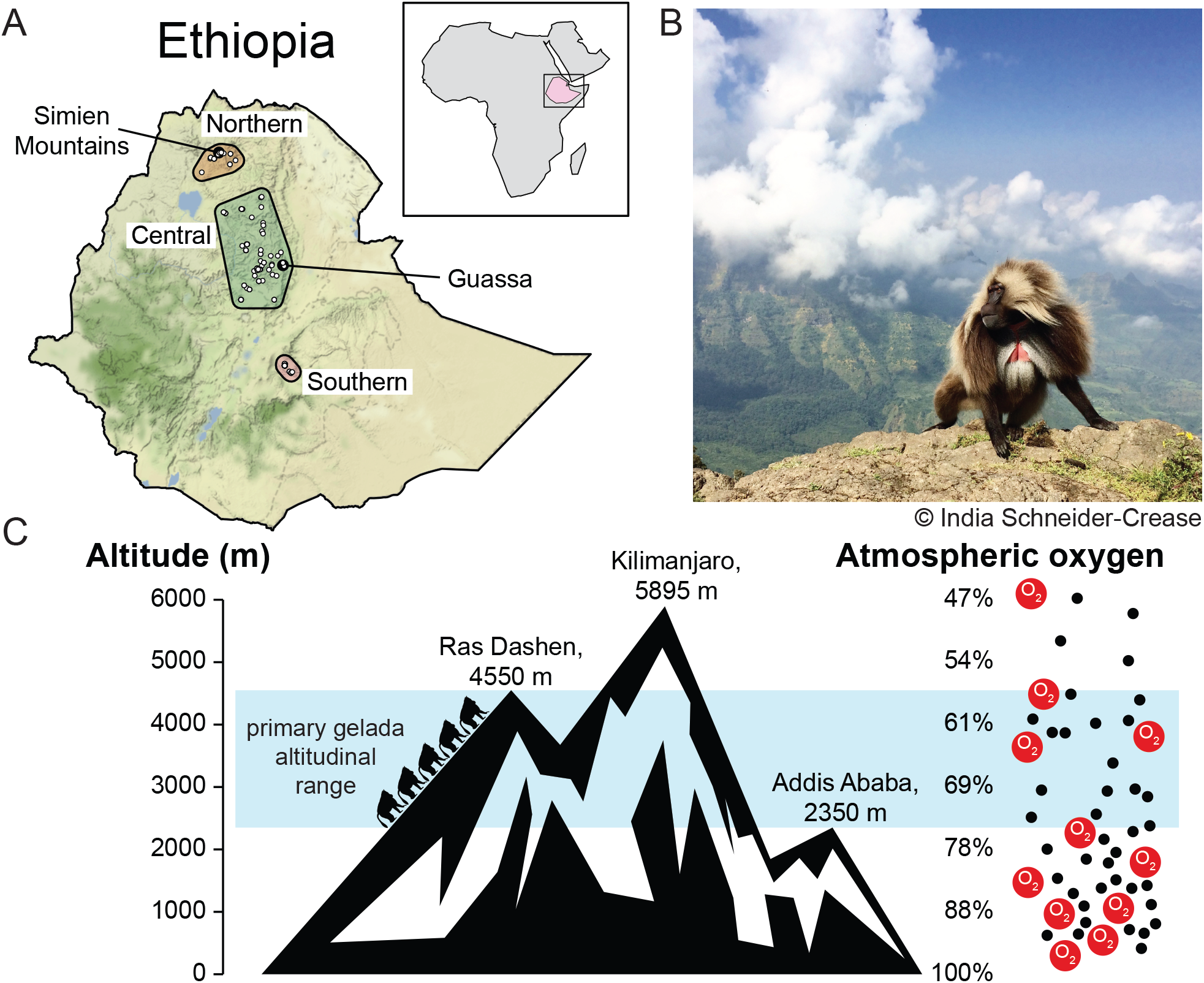
The gelada at high altitude. (A) Geladas form three main populations that are each geographically restricted to highland areas of Ethiopia. Presence points are shown from the sample of Zinner *et al.* [6]. (B) An adult male gelada in the Simien Mountains (photo © India Schneider-Crease). (C) Geladas are found almost exclusively from 2,350 to 4,550 meters above sea level, constituting one of the highest altitudinal ranges of any primate species.

Perspectives on gelada high-altitude adaptations are particularly important given their close evolutionary affinity and shared biology with humans and may lend insights into treatments for diseases and disorders associated with high altitude [14], including acute mountain sickness, high-altitude cerebral edema, and high-altitude pulmonary edema in high-altitude travelers, as well as chronic mountain sickness and preeclampsia in high-altitude residents. Furthermore, given the roles of hypoxia and ischemia in low-altitude diseases [15], a deeper understanding of mechanisms conferring resilience to hypoxia has the potential to inform treatment of diseases at all altitudes [16].

We sequenced and assembled the first gelada reference genome and combined it with detailed physiological, demographic, and morphological data collected from wild geladas to identify adaptations to their high-altitude environment. Through comparison with other mammals, we identified unique genomic adaptations to high altitude. We also analyzed population resequencing data from 70 wild and captive geladas to infer gelada population structure and demographic history. Curiously, we identified a chromosomal fission event that is polymorphic across at least two gelada populations and may act as a barrier to hybridization between populations, underscoring a potential case of incipient speciation in primates with important conservation implications.

## Sequencing, synteny, and annotation

We sequenced and assembled the genome of a wild adult female gelada from the Simien Mountains, Ethiopia, using a combination of two technologies: the linked-read 10x Genomics Chromium system [17] and Hi-C [18, 19]. Initial assembly of the 10x linked-read data (55.7-fold coverage) yielded a highly intact assembly (contig N50: 134.4 Kb; scaffold N50: 57.3 Mb), which was substantially improved by incorporating Hi-C intrachromosomal contact data to produce a reference assembly with chromosome-length scaffolds and comparable contiguity and coverage to other recent nonhuman primate genomes (contig N50: 310.1 Kb; scaffold N50: 130.2 Mb; Supplementary Fig. 1a). A BUSCO analysis of genome completeness identified 12,267 genes—of which 12,098 are one-to-one orthologs—comprising 91.7% and 89.0% of expected genes present and complete in mammals and primates, respectively [20] (Supplementary Fig. 1c). In total, our assembly includes 20,683 protein-coding genes annotated by NCBI [21]. The gelada genome is highly syntenic with closely related genomes, showing strong collinearity with the anubis baboon (*Papio anubis*) genome (Panu 3.0 [22]) (Supplementary Fig. 1b).

## Novel centric fission and incipient speciation

In assembling the genome of our reference individual, we identified an unexpected karyotype, 2n=44, that was not present in any other species in the papionin clade (macaques, drills/mandrills, mangabeys, baboons), which dates back to approximately 12 mya [26] and otherwise exhibits a conserved count of 21 chromosome pairs [23] (Fig. 2a). Our reference individual was homozygous for a centric fission [27] of chromosome 7, resulting in two new acrocentric chromosomes that we refer to as 7a and 7b (Fig. 2b,c). A single case of an apparently identical variant was previously reported in a captive gelada individual who was heterozygous for this variant (2n=43) but was interpreted as a rare structural anomaly in papionins [28]. We confirmed the homozygous 2n=44 karyotype via G-banding of fibroblasts in our reference individual and 3 additional unrelated individuals from the northernmost gelada population (Fig. 2c), demonstrating instead that this fissioned chromosome is a stable, possibly fixed variant in the Northern population. This variant appears to be unique to Northern geladas, with 2/2 wild geladas from central Ethiopia and 7/9 captive geladas from zoos—mainly of Central origin (Supplementary Fig. 7)—exhibiting the ancestral karyotype of 2n=42 (Fig. 2c and Supplementary Fig. 3). Two zoo geladas were heterozygous (2n=43), indicating a recent ancestor with a homozygous 2n=44 karyotype. These two heterozygous individuals had the most Northern ancestry (>10%) and the only Northern mitochondrial haplotype of all captive samples, suggesting that Northern ancestry can be traced through their maternal line (Fig. 2d and Supplementary Fig. 7). Together, the evidence is consistent with our hypothesis that the fissioned chromosome is a uniquely Northern trait. Despite having opportunity, neither individual successfully reproduced in captivity, suggesting the heterozygous karyotype may be associated with reproductive incompatibilities in hybrids, as is the case with other balanced chromosomal polymorphisms [29]. This chromosomal variant thus represents a possible barrier to hybridization that may underlie speciation between Northern and Central geladas. These groups are typically considered subspecies—*T. gelada gelada* (Northern) and *T. gelada obscurus* (Central)—but show evidence of being distinct evolutionary units that would qualify as species under the phylogenetic species concept [30]. If the centric fission is fixed or near-fixation in Northern geladas and the heterozygous karyotype is associated with reduced fitness, as our data suggest, these populations would further qualify as species under the biological species concept, cementing the case for taxonomic revision and reconsideration of conservation priorities. Furthermore, this would provide a possible case study of chromosomal rearrangements underlying speciation, the mechanisms of which remain poorly understood [31, 32].

**Figure 2.**
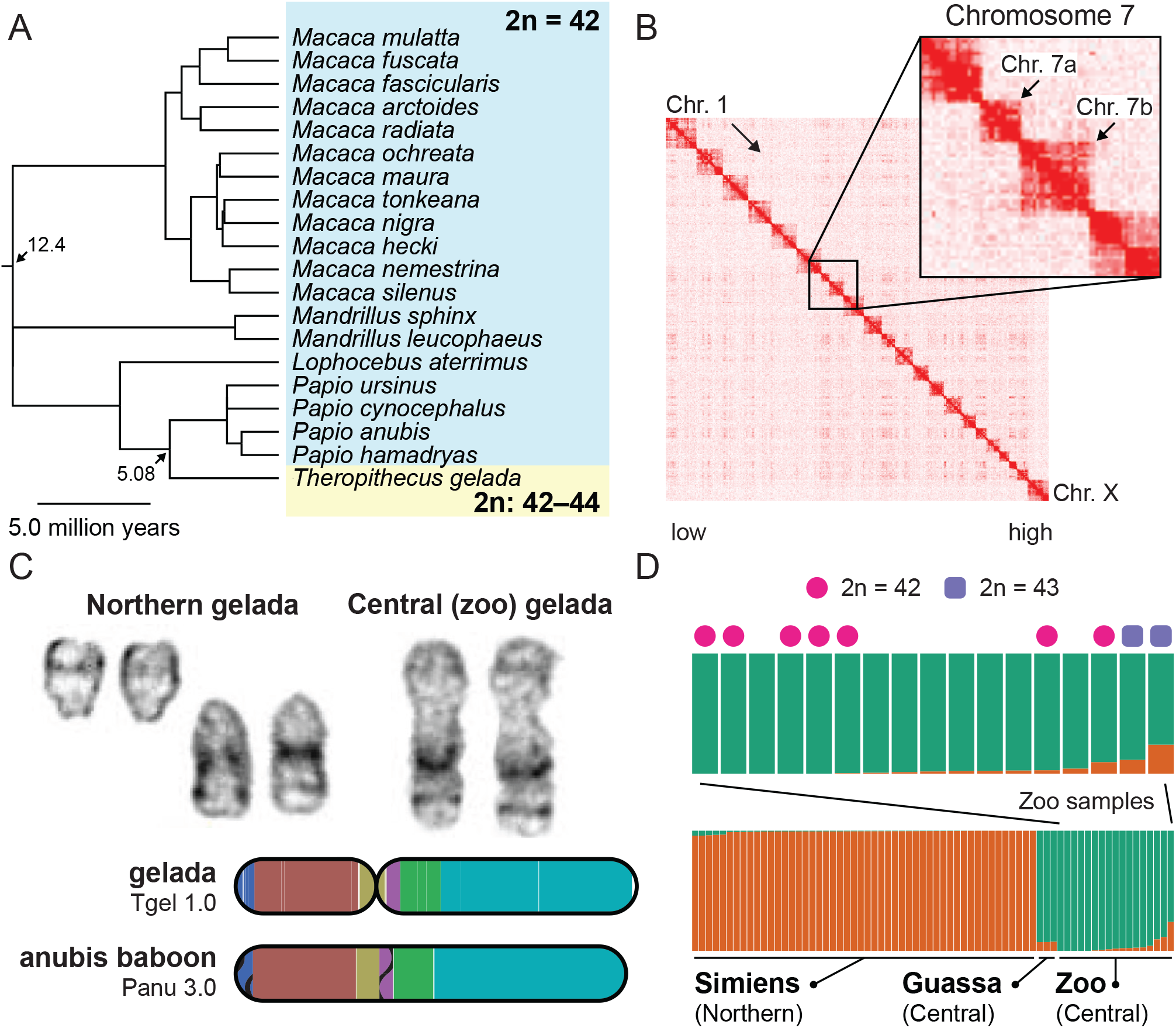
Unique karyotypic evolution in geladas. (A) Apart from geladas, the papionin clade exhibits an extremely conserved karyotype of 42 diploid chromosomes. 20 species with known karyotypes sampled by Stanyon *et al.* [23] are shown with the consensus chronogram from TimeTree [24, 25]. (B) Hi-C contact map reveals a distinct lack of contacts between the arms of chromosome 7. (C) G-banded karyotyping and analysis of genomic rearrangements reveal strong synteny between fissioned chromosomes and the intact arms of chromosome 7 in Central geladas and baboons, respectively. (D) Population structure of 70 resequenced gelada genomes reveals two main populations differentiating Northern (orange) and Central (green) geladas. Zoo animals are of mainly Central ancestry, but two individuals with the highest levels of Northern ancestry are also heterozygous for the centric fission characteristic of Northern geladas.

## Conservation and population genomics

To better understand the demographic history of geladas, including historical population sizes and population divergence, we sequenced the whole genomes of 70 captive and wild geladas from multiple parts of Ethiopia (n=3 wild Central geladas; n=50 wild Northern geladas; n=17 captive geladas of Central origin) as well as 20 hamadryas baboons from Filoha, Ethiopia [33] (median coverage = 11.5x; Supplementary Table 1). Our sample did not include any individuals from the Southern gelada population, which is difficult to access but represents another distinct evolutionary unit [30] (Fig. 1a). We used the Multiple Sequentially Markovian Coalescent (MSMC) to infer the demographic history of sequenced geladas, which indicated that the effective population sizes of the two gelada populations (Northern and Central) began to diverge about 500 thousand years ago (Fig. 3a). It is therefore most likely that the chromosomal fission arose in Northern geladas following this population divergence.

**Figure 3.**
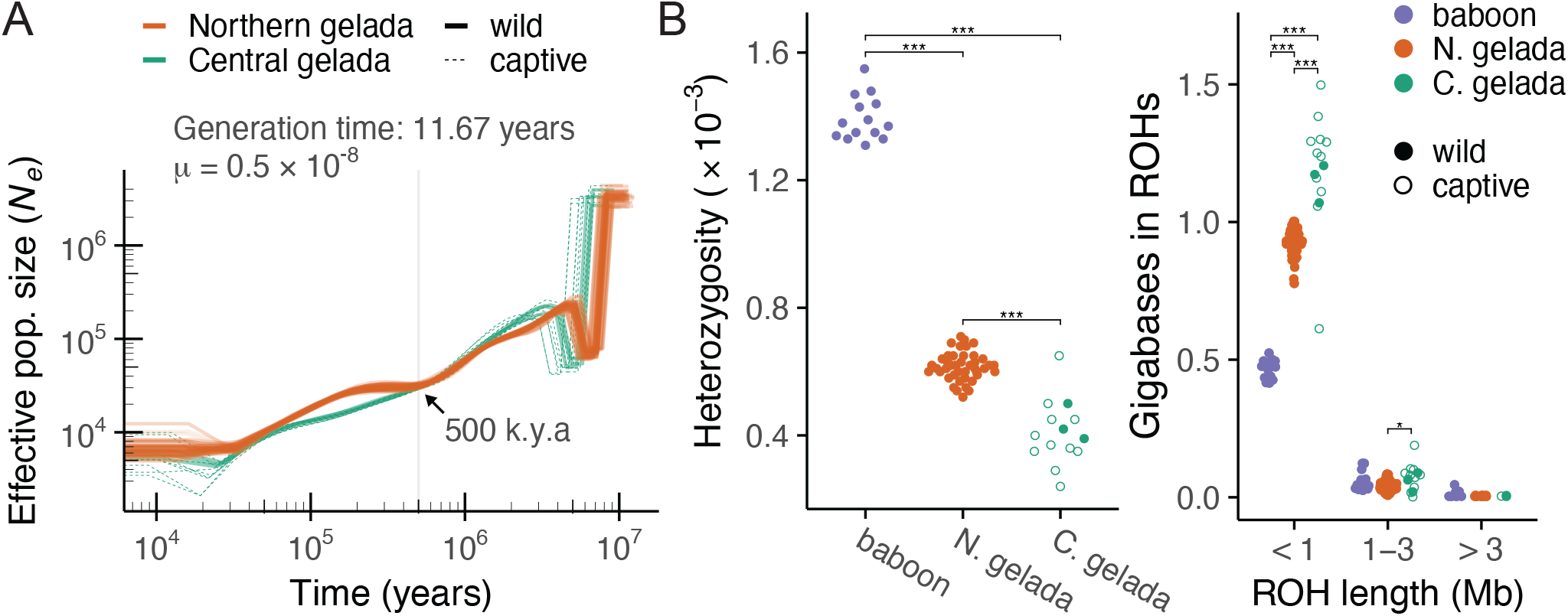
Historical demography and genomic diversity among gelada populations. (A) Multiple Sequentially Markovian Coalescent model reveals a historical divergence in effective population size between Northern and Central geladas, occurring roughly 500 k.y.a. (B) Analysis of genomic diversity reveals that geladas have lower heterozygosity and a higher portion of the genome in runs of homozygosity (ROHs) relative to hamadryas baboons, indicating less genetic diversity and a lower effective population size. Within geladas, the Northern population is more diverse than the Central population according to both metrics.

The geladas in our sample fell into two distinct populations corresponding to previously described subspecies: the Northern population, which encompasses all wild individuals from the Simien Mountains, and the Central population, which encompasses wild individuals from Guassa Community Conservation Area as well as the majority of individuals from zoos (Fig. 2d; based on unsupervised clustering [34]). A small number (n = 3) of zoo individuals showed elevated (>9%) fractions of Northern ancestry, including the two zoo animals found to have 2n=43 heterozygous karyotypes. These cases are likely explained by captive breeding of parents from different populations. We found no evidence of interbreeding between wild gelada populations.

We found higher genetic diversity in the Northern gelada population than the Central gelada population (Fig. 3b). Central geladas had lower median heterozygosity than Northern geladas (Wilcoxon, *P* = 8.7e-07) and also significantly longer runs of homozygosity, specifically for runs <1 Mb (Wilcoxon, *P* = 8.5e-08) and 1–3 Mb (*P* = 0.008). Both gelada populations were significantly less genetically diverse than hamadryas baboons (median heterozygosity: *P* = 2.6e-08 [Northern], *P* = 7.4e-06 [Central]), perhaps reflecting their limited geographic distribution and habitat discontinuity compared to baboons.

## Physiological adaptations to high-altitude hypoxia

### Hemoglobin-oxygen affinity

Many animals that have adapted to high altitude have evolved an increased affinity of oxygen to hemoglobin, which can minimize the decline in arterial oxygen saturation in spite of environmental hypoxia [36]. We therefore examined the genes encoding the alpha- and beta-chain subunits of adult hemoglobin in geladas. We found two amino acid substitutions in hemoglobin-alpha, at sites 12 and 23, that are unique to geladas relative to other primates (Fig. 4a). To test whether these substitutions alter functional properties of the protein, we measured hemoglobin-oxygen binding affinity from purified adult hemoglobin of geladas, humans, and three species of baboons (Supplementary Table 8). In the presence of allosteric cofactors—the experimental condition most relevant to *in vivo* conditions—we found no differences in *P*_50_) (the partial pressure at which hemoglobin is 50% saturated) of gelada hemoglobin compared to that of humans (*P* = 0.053) or baboons (*P* = 0.950) (Fig. 4b). Thus, the amino acid substitutions found in gelada hemoglobins do not appear to be associated with increased hemoglobin-oxygen affinity, in contrast with the pattern generally observed in high-altitude birds [36] and some high-altitude mammals [4, 37, 38] but mirroring a similar lack of increased oxygen affinity in snow leopard hemoglobin [39].

**Figure 4.**
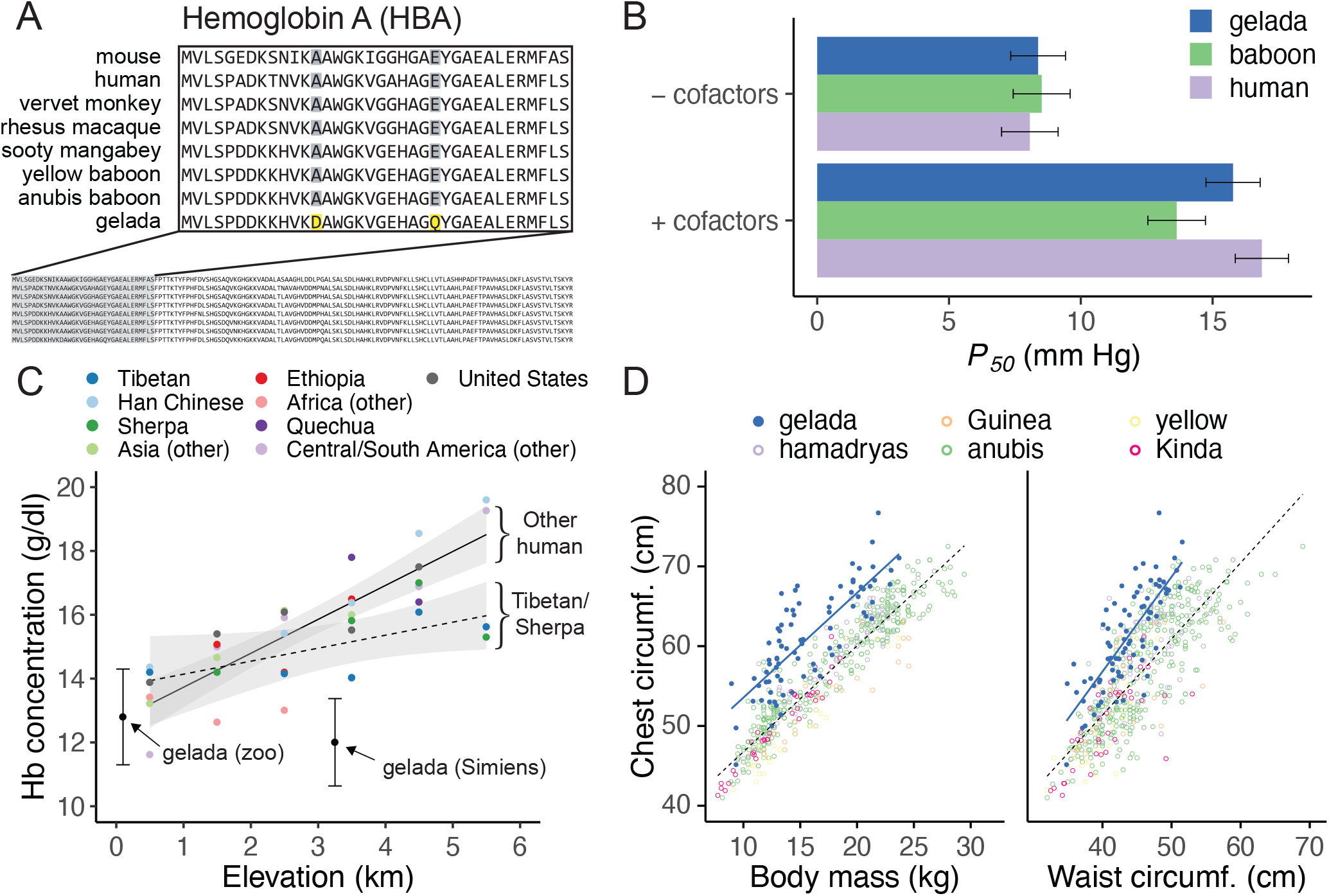
Gelada blood and lung phenotypes at high altitude. (A) Protein alignment reveals two unique substitutions in the alpha subunit of hemoglobin (Hb) in gelada. (B) Hb–O_2_ affinity assays, however, do not find evidence of increased oxygen binding affinity (i.e., lower *P*_50_) of gelada Hb. (C) Geladas at high altitude do not exhibit elevated Hb concentrations (erythrocytosis) at high altitude, in contrast to most humans with the notable exception of Tibetans and Sherpa. Values for human populations are plotted from the metaanalysis by Gassmann *et al.* [35]. The mean ± standard deviation is shown for zoo and wild (Simiens) geladas. (D) Comparison of gelada chest circumferences to those of five baboon species reveals that geladas maintain larger chest circumferences relative to their body mass and waist circumference, respectively.

### Hemoglobin concentration and erythrocytosis

In lowland mammals, a typical response to chronic hypoxia is an increase in red blood cell production (erythrocytosis), which in humans is associated with acute mountain sickness in high-altitude travelers and chronic mountain sickness in high-altitude residents. Erythrocytosis is particularly prevalent in Andean highlanders and is altitude-dependent in its severity [40, 41], while hemoglobin concentrations notably remain low in Tibetans [42]. To test for erythrocytosis in geladas, we compared hemoglobin concentrations from 92 wild geladas sampled at high altitude (3250–3600 m) to values reported from captive geladas [43] and baboons [44] at low altitude. We found that hemoglobin concentrations in geladas at high altitude were not elevated and were in fact significantly lower than hemoglobin concentrations in either captive geladas (*P* = 0.005) or baboons (*P* < 0.001). The absence of elevated hemoglobin concentrations in wild geladas living at high-altitude is consistent with patterns documented in other hypoxia-adapted alpine mammals [45, 46] and, among humans, most closely resembles the Tibetan phenotype (Fig. 4c). Since red blood cell production is induced by reduced oxygenation of renal tissue, the absence of an elevated hemoglobin concentration in wild geladas living at >3000 m suggests that the animals are able to sustain adequate tissue-oxygen delivery in spite of the reduced availability of oxygen. Such physiological compensation may be attributable to evolved and/or plastic changes in any number of cardiorespiratory or circulatory traits that govern oxygen-transport.

### Pulmonary adaptations at high altitude

We also tested the hypothesis that geladas might compensate for hypoxia by expanding their lung volumes, which is a known high-altitude developmental adaptation spurred by rapid lung growth in early life [47]. Expanded lungs in high-altitude animals maximize the pulmonary diffusing capacity for oxygen by proliferating alveolar units and increasing surface area for gas exchange [48, 49]. To test for correlates of expanded lung volumes in geladas, we compared chest circumferences in wild geladas (n=78) to an extensive database of baboon morphometric measurements (n=482) [50, 51, 52]. We found that, controlling for sex and baboon species, geladas had significantly larger chest circumferences compared to baboons when also controlling for body mass (*P* = 1.63e-42), waist circumference (*P* = 3.23e-31), or both (*P* = 8.58e-46) (Fig. 4d). These results indicate that geladas have larger relative chest circumferences and thus potentially expanded lung capacity, which parallels the larger chest dimensions exhibited by native Andean highlanders [53]. It is currently unknown whether these differences have a genetic basis.

## Genomic adaptations to high altitude

To identify signatures of adaptations to high altitude in the gelada genome, we first focused on two forms of genetic change that could underlie changes in phenotype: coding mutations that alter protein function and gene duplications that alter gene dosage and/or division of labor among protein isoforms.

We tested for evidence of positive selection in gelada coding sequences using two complementary *d*_*N*_/*d*_*S*_ tests—the site-based model implemented in **PAML** [54] and the gene-based model implemented in **BUSTED** [55]. We assigned proteins from 40 taxa (Supplementary Fig. 4 and Supplementary Table 2) to single-copy orthogroups [56] and, after filtering (Methods), included 6,105 protein-coding genes in our analysis. We identified 103 genes exhibiting significant signatures of positive selection (FDR-adjusted *P* < 0.05) using both *d*_*N*_/*d*_*S*_ approaches.

To test for gene duplication resulting in significant expansions of gene families, we assigned proteins from 40 taxa (Supplementary Fig. 4 and Supplementary Table 2) to gene families in the TreeFam9 database [57, 58]. We then tested for gene family size changes using birthdeath models implemented in **CAFE** [59]. We identified 108 gene families exhibiting significant expansions in gene family size (FDR-adjusted *P* < 0.05).

### Positive selection on protein-coding sequences

We found several compelling candidate genes for high-altitude adaptation among 103 total genes with significant signatures of positive selection (FDR-adjusted *P* < 0.05; Supplementary Table 3). These included four genes involved in the hypoxia-inducible factor (HIF) pathway (*ITGA2*, *NOTCH4*, *FERMT1*, *MLPH*). We also identified several that have been identified as candidate genes in human hypoxia-adapted populations, including *FRAS1*, which is involved in renal agenesis and exhibits signatures of positive selection in Tibetans [60] and Ethiopians [61], *HMBS*, which is involved in heme biosynthesis and exhibits a signature of positive selection in Nepalese Sherpa [62], and *TNRC18*, a largely unknown gene that is linked to selection in Bajau breath-hold divers [63]. Other notable candidate genes include *AQP1*, which plays an important role in fluid clearance and edema formation following acute lung injury [64], *COX15*, which is involved in heme *a* biosynthesis and cytochrome c oxidase assembly [65] and exhibits signatures of positive selection in high-altitude rhesus macaques [66], *DHCR24*, which is involved in the induction of heme oxygenase 1 (HO-1) [67] and exhibits a signature of positive selection in alpine sheep [68], and *CYGB*, which is part of the globin family and encodes an oxygen-binding respiratory protein [69].

At the pathway level, we found that signatures of positive selection were enriched (FDR-adjusted *P* < 0.1; Supplementary Table 4) for processes related to classical functions associated with high-altitude adaptation, including oxygen sensing (response to hypoxia; angiogenesis; cellular response to hypoxia) [70], response to oxidative (response to hydrogen peroxide) and other stress (response to glucocorticoid; MAPK cascade) [71, 72], and female reproduction (in utero embryonic development; response to estradiol; ovulation cycle process; female pregnancy) [73, 74, 75]. In addition, we identified several enriched processes related to neural function (axon guidance; positive regulation of neuron projection development; chemical synaptic transmission; brain development), cell growth and proliferation (response to insulin; negative regulation of cell population proliferation; negative regulation of canonical Wnt signaling pathway), and cardiac function (cell-cell signaling involved in cardiac conduction).

While we found a high degree of overlap between putatively selected pathways in geladas and human populations living at high altitudes, aside from notable examples listed above, few candidate genes identified by our analysis were shared with candidate genes identified by studies of high-altitude human populations [2, 16] or other high-altitude primates [13, 66]. This suggests that gelada adaptations to similar physiological challenges at high altitude largely involve different suites of genes [76], underscoring their utility as a novel model for understanding adaptations to hypoxia.

### Gene family expansion

We identified 108 gene families with significant expansion in the gelada lineage compared to 43 gene families with significant contractions (FDR-adjusted *P* < 0.05; Supplementary Table 5). Significant expansions included the genes *CENPF* and *SART1*, which each exist as a single copy in most primates, including the common ancestor of geladas and baboons, but have expanded to include five copies and four copies respectively in the gelada lineage (Supplementary Fig. 5b,c). *CENPF* encodes a kinetochore protein that regulates chromosome alignment and separation during mitosis and also protects centromeric cohesion [77]. Interestingly, *CENPF* is a marker for cell proliferation in human malignancies [78] and is strongly upregulated in response to hypoxia in bone marrow mesenchymal cell cultures [79], suggesting that it may also play a role in the response to high-altitude hypoxia.*SART1* suppresses activation of HIF-1 by promoting the ubiquitination of HIF-1*a*. Expansion of the*SART1* family may therefore be a possible adaptation for suppressing constitutive HIF-1 activation under conditions of chronic environmental hypoxia.

We also identified biological processes that were associated with signatures of gene family expansion (FDR-adjusted *P* < 0.1; Supplementary Table 6). We found that signatures of gene expansion were significantly associated with processes related to the hypoxia response (regulation of transcription from RNA polymerase II promoter in response to hypoxia) as well as the DNA damage response (e.g., DNA repair; nucleotide-excision repair; DNA incision, 5’-to lesion; DNA duplex unwinding), which may reflect degrees of DNA damage due to elevated levels of ultraviolet radiation at high altitude [80, 81]. Other enriched processes included those related to immune function (e.g., NIK/NF-kappaB signaling; stimulatory C-type lectin receptor signaling pathway; viral transcription; IL-1-mediated signaling pathway; T cell receptor signaling pathway; TNF-mediated signaling pathway), cell proliferation (e.g., Wnt signaling pathway; planar cell polarity pathway; MAPK cascade), oxidative phosphorylation (mitochondrial respiratory chain complex I assembly; mitochondrial electron transport, NADH to ubiquinone; oxidation-reduction process), and hematopoiesis (regulation of hematopoietic stem cell differentiation).

### Accelerated evolution in the gelada lineage

To investigate the emergence of gelada-specific features and to expand our analysis to non-coding regions of the genome including regulatory elements [82], we identified and characterized genomic regions that are highly conserved through evolution but exhibit a greater number of changes in the gelada lineage. A similar approach has been used to identify “human accelerated regions” (HARs) that are possible hallmarks of human evolution [83, 84], tend to be developmental gene regulatory elements or in non-coding RNA regions [85], and are putatively linked to uniquely human social behavior and cognition [86].

We used an approach modeled on that of Pollard *et al.* [84] to define uniquely accelerated regions in the gelada lineage, which we refer to as “gelada accelerated regions”, or GARs. We analyzed 60,345 conserved alignment blocks across a total of 57 mammalian taxa (Methods), including geladas, and identified a total of 29 GARs (FDR-adjusted *P* < 0.2; Supplementary Fig. 6 and Supplementary Table 9). We identified fewer GARs than reported counts of HARs, which range from approximately 200–3000 at similar thresholds [84, 87], likely due to differences in filtering, thresholding, and other aspects of methodology.

Of the 29 GARs, 13 (44.9%) were located in intergenic regions, ten (34.5%) in introns, one (3.4%) in a 5’ UTR region, and the remaining five (17.2%) in coding sequences. Many of these GARs were nominally regulatory: 13 GARs (44.9%) were associated with regulatory hallmarks of enhancer activity in at least one primary tissue or cell type in humans (Methods). Of these putative enhancers, 11 are associated with hallmarks of enhancer activity in human fetal tissues or were nearest to genes that are involved in developmental processes including in utero embryonic development and post-embryonic development. These results indicate that a large fraction of GARs may function as developmental enhancers, similar to HARs [87]. Two additional GARs (GAR5 and GAR8) are located in regions showing strong evidence of being transcriptional start sites across many tissues (> 60 cell/tissue types each).

Strikingly, in two cases, multiple GARs were found near the same genes. These genes were *RBFOX1*, which was the closest gene to GAR28 and GAR29, and *ZNF536*, which was the closest gene to GAR26 and GAR27. In both cases, GARs were at least 500 kb apart from one another and in low linkage disequilibrium (mean *r2*: 0.07–0.17). Both *RBFOX1* and *ZNF536* are linked to brain expression and function. *RBFOX1* is an important regulator of neuronal excitation [88] while *ZNF536* is a negative regulator of neuronal differentiation [89]. None of the associated regions showed hallmarks of transcription factor binding or chromatin accessibility in human tissues and cells, making their function at present a mystery.

Other identified GARs also were located nearest to genes involved in brain function. These genes included *RTN4RL1* (GAR21), which is involved in postnatal brain development and regulating regeneration of axons, *GIGYF2* (GAR16), a regulator of vesicular transport and IGF-1 signaling in the central nervous system [90], *CNTN4* (GAR3), which has been linked to neuropsychiatric disorders and fear conditioning [91], and *NFASC* (GAR1), which is linked to neurite outgrowth and adhesion [92]. The accelerated evolution of GARs near multiple genes related to neural function in geladas may reflect the sensitivity of the brain to the metabolic pressures of high-altitude hypoxia [93, 94].

Several GARs were located nearest to genes that are involved in the response to hypoxia or oxidative stress, suggesting that they might be adaptations to high altitude environments [72]. Two GARs were located nearest to genes—*HTATIP2* (GAR17) and a novel gene in geladas (ENSTGEG00000009621; GAR7)—involved in oxidation reduction. One GAR was located in the intron of *RCAN1* (GAR4), which is involved in the response to oxidative stress and regulation of angiogenesis [95]. Another GAR was located in the 5’ UTR region of *FBN1* (GAR8), which is hypoxia responsive [96] and more highly expressed at higher elevations among yaks [97].

Intriguingly, we found that one GAR, GAR18, was nearest to the gene *SOX6*, which plays an essential role in erythroid cell differentiation and is necessary for basal and stress erythropoiesis [98, 99]. GAR18 was found 2,651 bp upstream of *SOX6* and was associated with regulatory hallmarks of enhancer activity in five primary cell types (Supplementary Table 9). Given its position and putative function as an enhancer in humans, GAR18 could suppress hypoxia-induced erythropoiesis by decreasing or disabling enhancer activity, providing a direct link to the lack of altitude-associated erythropoiesis that we observed in wild geladas.

## Conclusion

The first assembled gelada genome provides novel insights into the unique adaptations of this charismatic Ethiopian primate. We identified a novel and stable karyotype that appears to be at extremely high frequency and possibly fixed in the Northern population of geladas. Given that chromosomal rearrangements tend to be associated with infertility in heterozygous karyotypes, our findings suggest that geladas may encompass at least two distinct biological species. This finding is important for at least two reasons. First, a taxonomic revision would roughly halve the populations of each gelada species and, consequently, alter their conservation status and ultimately increase resources to protect them. Second, the centric fission of chromosome 7 is an extraordinarily recent example of a stable chromosomal variant in a long-lived primate. It therefore provides a unique opportunity to study karyotypic evolution, the birth of new centromeres, and the role of chromosomal rearrangements in speciation in a primate closely related to humans.

By combining morphometric, hematological, and genomic data, we identified a suite of gelada-specific traits that may confer adaptation to their high-altitude environment, including evidence for increased lung capacity and positive selection in a number of hypoxia-related genes, and gelada-lineage-specific accelerated regulatory regions. Interestingly, while we found gelada-specific amino acid substitutions in hemoglobin, these changes did not alter oxygen-binding affinity, which highlights the need for functional assays to validate purely sequence-based findings. With this in mind, our genome assembly and gelada-specific genetic changes provide multiple avenues for future research on the function of the protein-coding and regulatory changes unique to geladas. This research thus builds upon our current understanding of the mechanisms of adaptation to extreme environments and provides an avenue for research that may have a transformative impact on the study and treatment of hypoxia-related conditions.

## Acknowledgments

We are grateful, first and foremost, to those who made this research possible, particularly the research staff (Esheti Jejaw, Ambaye Fenta, Setey Girmay, Dereje Bewket, and Atirsaw Adwana), logistical support staff (Tariku W/Aregay and Shiferaw Asrat), and assistants and students of the Simien Mountains Gelada Research Project, as well as the Ethiopian Wildlife Conservation Authority (EWCA) for permission and support for working in the Simien Mountains National Park. We are also grateful to EWCA, the Amhara Regional Government, and Mehal Meda Woreda for permission to conduct research at Guassa Community Conservation Area; and to Badiloo Muluyee, Ngadaso Subsebey, Bantilka Tessema, Tasso Wudimagegn, and many field assistants for important logistical research support there. We thank David McDonald and the Cellular Imaging Core at the Fred Hutchinson Cancer Research Center for assistance with karyotyping. We are additionally grateful to Sierra Sams and Sarah Ford for assistance with lab work, and to Michael Montague, Kelley Harris, Abigail Bigham, Graham Scott, Ivan Liachko, Zev Kronenberg, Olga Dudchenko, Noah Simons, Nelson Ting, and Julien Dutheil for feedback through various stages of this research.

Support for this research was provided by the National Science Foundation (BCS 2010309, BCS 1848900, BCS 2013888, BCS 1723237, BCS 1723228, BCS 0715179, OIA 1736249, IOS 2114465, IOS 1255974, and IOS 1854359), the National Institutes of Health (NIA R00AG051764 and NHLBI R01HL087216), the University of Washington Royalty Research Fund, the San Diego Zoo, and the German Research Foundation (DFG KN1097/3-1). KLC was supported by a National Institutes of Health fellowship (NIA T32AG000057). MCJ was supported by the Natural Environment Research Council (NE/T000341/1) and the Natural Sciences and Engineering Research Council Discovery Accelerator Grant. ISC (Schneider-Crease) is supported by the ASU Center for Evolution and Medicine.

## Author contributions

NSM, KLC, and MCJ conceived the research. KLC, MCJ, ISC (Schneider-Crease), ADM, AL (Lu), JCB, TJB, and NSM designed the study. KLC, ISC (Schneider-Crease), SS, FA, ISC (Chuma), SK, AL (Lemma), BA, JCB, TJB, and NSM collected field gelada samples and data, facilitated by AAH and FK. PJF, NN, CM, MLH, JDW, ASB, CMB, JR, JEPC, and CJJ contributed samples and/or data. AVS and JFS designed, performed, and analyzed Hb-O2 affinity experiments. KLC, AMD, and NSM generated genomic data. KLC, MCJ, and NSM performed genomic analyses. KLC, MCJ, and NSM wrote the paper. All authors revised and approved the final manuscript.

## Data availability

All genomic data, including the Tgel 1.0 assembly (GenBank accession number GCA_003255815.1) and short-read sequencing data, are available through National Center for Biotechnology Information (NCBI) repositories and are linked to BioProject accession number PRJNA470999. Gelada hematological and morphological data are available on Dryad (https://doi.org/10.5061/dryad.fbg79cnvq).

## Online Methods

### Animal procedures

#### Capture and release

Samples and data collected for this study were obtained from wild geladas in the Simien Mountains National Park (~3,000–4,550 meters above sea level) as part of continuous long-term research conducted by the Simien Mountains Gelada Research Project (SMGRP). Beginning in 2017, the SMGRP has carried out annual capture-and-release campaigns during which animals were temporarily immobilized through remote-distance injection. Briefly, a mixture of ketamine (7.5 mg/kg) and medetomidine (0.04 mg/kg) was injected using darts delivered by a blowpipe (Telinject USA, Inc). Following data and sample collection under the supervision of licensed veterinarians and veterinary technicians, immobilization was reversed with atipamezole (0.2 mg/kg). Animals were monitored by project staff throughout their recovery until they were visibly unimpaired and had returned to their social units. All research was conducted with permission of the Ethiopian Wildlife and Conservation Authority (EWCA) following all laws and guidelines in Ethiopia. Animal procedures were conducted with approval by the Institutional Animal Care and Use Committees (IACUCs) of the University of Washington (protocol 4416-01) and Arizona State University (20-1754R). This research conformed to the American Society of Primatologists/International Primatological Society Code of Best Practices for Field Primatology.

#### Morphometrics

While animals were sedated, we collected morphometric measurements including body mass, chest circumference, and waist circumference. Body mass was measured by a hanging digital scale to 0.05 kg precision. Chest circumference and waist circumference were measured using flexible tape to 0.1 cm precision. Chest circumference was defined as the maximum circumference of the trunk, taken at the maximum anterior projection of the thoracic cage. Waist circumference was defined as the minimum circumference between the pelvis and the thoracic cage.

#### Biological sample collection

Whole blood was obtained from all chemically immobilized individuals by femoral venipuncture and collected into K3 EDTA S-Monovette collection tubes (Sarstedt). 1 ml of whole blood was cryopreserved in liquid nitrogen, ~50 μl was used for hematology, and the remainder was fractionated by centrifugation using a ficoll gradient. Fibroblasts were cultured from small biopsy punches of ear tissue that were stored in RPMI supplemented with 20% FBS and 10% DMSO. To maximize viability, these samples were frozen in steps by first storing in styrofoam at −20°C overnight, then transferring to liquid nitrogen.

We also measured hemoglobin concentrations using an AimStrip 78200 digital hemoglobin meter. We loaded 10 μl of venous blood into provided test strips and recorded hemoglobin concentrations (g/dl) using the digital meter.

#### Other DNA sources

Apart from primary DNA samples collected for this project (n=50), we obtained additional DNA samples from other sources, including DNA extracts from 20 wild hamadryas baboons from Filoha, Ethiopia (contributed by C. Jolly and J. Phillips-Conroy), which were previously determined to have unadmixed ancestry [33], DNA extracts from 17 zoo geladas (n=1 contributed by the Wildlife Conservation Society/Bronx Zoo, n=16 contributed by the San Diego Zoo Wildlife Alliance), and muscle tissue from 3 wild geladas (contributed by N. Nguyen and P. Fashing). A full list of DNA samples used for this research is provided in Supplementary Table 1.

### Sample collection, sequencing, and assembly

#### 10x Genomics Chromium library generation and sequencing

High molecular weight DNA was extracted from cryopreserved whole blood of an adult eight year-old female gelada (DIX) from the Simien Mountains using the Gentra Puregene Blood Kit (Qiagen) following manufacturer instructions and quality-checked using pulsed-field gel electrophoresis. Linked-read libraries were then prepared using the Chromium Genome Reagent Kit v2 (10x Genomics) following manufacturer instructions. Finished libraries were sequenced to 55.7x coverage on two lanes of the Illumina HiSeq X platform using 2×150 bp sequencing.

#### Hi-C library generation and sequencing

Approximately seven million peripheral blood mononuclear cells (PBMCs) were isolated by ficoll gradient, washed, counted, fixed in formalin, and cryopreserved. Hi-C libraries were later prepared from cryopreserved formalin-fixed PBMCs following Rao *et al.* [18] and sequenced on the Illumina NextSeq 500 platform using 2⇥81 bp sequencing.

#### Genome assembly

Chromium-derived reads were assembled using `Supernova` v2.0.1 [17] with default parameters. Resulting scaffolds were then further assembled incorporating Hi-C data through the 3D *de novo* assembly (`3D-DNA`) pipeline v170123 [19]. Hi-C contact maps and the draft assembly with chromosome-length scaffolds were edited using `Juicebox Assembly Tools` [100] to correct visually apparent misjoins. Finally, gaps were closed using `GapCloser` v1.12 [101] to produce the final assembly (Tgel 1.0).

### Transcriptome sequencing and genome annotation

RNA was obtained from cultured fibroblast cell lines derived from a biopsy ear punch taken from the same adult female gelada (DIX) and an unrelated male gelada (DRT_2017_018) from the Simien Mountains National Park, Ethiopia. Total RNA was extracted using the Quick RNA Miniprep Plus kit (Zymo Research). RNA-seq libraries were prepared using the NEBNext Ultra II RNA Library Prep kit (New England Biolabs) following instructions for a 200 bp insert. Finished libraries were sequenced on the Illumina NextSeq 500 platform using 2×81 bp sequencing.

The final assembly (Tgel 1.0) was deposited in GenBank (accession GCA_003255815.1) and annotated de novo by the National Center for Biotechnology Information (NCBI) using the Eukaryotic Genome Annotation Pipeline [21]. Fibroblast RNA-seq short reads were submitted to the Sequence Read Archive (accessions SRX4071585 and SRX4100999) and included in the annotation pipeline.

### BUSCO assessment

To assess the completeness of the Tgel 1.0 genome assembly, we used `BUSCO` v4.0.6 [102] and compared our assembly against common orthologs in both the mammalian (dataset mammalia_odb10, creation date 2019-11-20) and primate (dataset primates_odb10, 2019-11-20) lineages (Supplementary Fig. 1c). `BUSCO` was run with default settings, using the following versions of third party components: `python` v3.7.4, NCBI `BLAST` v2.2.31, `Augustus` v3.2.3, `HMMER` v3.1b2, `SEPP` v4.3.10, and `Prodigal` v2.6.3.

### Synteny analysis

We assessed the synteny between chromosomal scaffolds of Tgel 1.0 and the anubis baboon reference genome, Panu 3.0 [22], using two approaches. In the first approach, we computed pairwise alignments between genomes using `nucmer` from `MUMmer` v3.23 [103], using a cluster size of 400, a minimum match length of 10 bp, and a maximum of 500 bp between clusters. We then used the `delta-filter` utility program from `MUMmer` to retain only alignments with a minimum identity of 40% and a minimum overlap of 1% between query and reference alignments. We then plotted links between assemblies (Supplementary Fig. 1b). In the second approach, we used the reference-free method implemented in `Smash` v1.0 [104] to identify syntenic blocks and to visualize chromosome rearrangements (Supplementary Fig. 2). We ran `Smash` using default settings.

### Karyotype assessment

Our Hi-C data revealed a distinct lack of contacts between scaffolds corresponding to nonoverlapping segments of chromosome 7 on the baboon reference genome (Fig. 2b). To test for a possible chromosomal fission in our reference individual, we performed G-banded karyotyping on fibroblasts cultured from the same individual, which confirmed that our reference had a homozygous karyotype with a centromeric fission in chromosome 7 (Fig 2c and Supplementary Fig. 3), resulting in two new acrocentric chromosomes that we refer to as 7a and 7b and a full karyotype of 2n=44. We tested for the presence of this centric fission in additional captive and wild geladas from zoos, northern Ethiopia (Simien Mountains), and central Ethiopia (Guassa). We counted chromosomes for zoo and northern Ethiopian geladas using karyotyping (with or without chromosome banding), taking advantage of the availability of live cells either through the Frozen Zoo (San Diego Zoo Wildlife Alliance) or through samples collected by our project.

For wild central Ethiopian geladas, for which live cells were not available, we tested for the presence of a chromosomal fission by generating and analyzing Hi-C data from ethanol-fixed tissue samples. We generated Hi-C libraries using the Proximo Hi-C animal kit (Phase Genomics) following manufacturer instructions and sequenced them on the Illumina iSeq platform. We then ran the `3D-DNA` pipeline v170123 [19] separately from each Hi-C library using our reference gelada chromosomal scaffolds as input. Resulting contact maps were then assessed for the presence/absence of contacts between 7a and 7b, both visually (Supplementary Fig. 3c) and by permutation. For permutations, we simulated the null distribution of interchromosomal contacts (i.e., contacts between distinct chromosomes excluding chromosome 7) by dividing the reference genome into 10 million base-pair windows, then randomly sampling windows without replacement until the combined sizes added up to the lengths of 7a and 7b, respectively. We then determined the frequency of Hi-C contacts between windows assigned to these simulated “chromosomes”. In all cases, we determined significant overrepresentation of contacts between arms of chromosome 7 relative to our simulated null distributions, thus rejecting the hypothesis that chromosome 7 is exclusively fissioned (i.e., 2n=44). Because the relative proportion of contacts between 7a and 7b surpassed estimates generated from baboon Hi-C data, we also rejected the possibility of a heterozygous karyotype (i.e., 2n=43), as baboons—along with all non-gelada papionins—are not known to exhibit a fissioned chromosome 7.

### Hemoglobin-oxygen (Hb-O_2_) affinity

To identify unique amino acid substitutions in geladas, we used amino acid sequences for the hemoglobin alpha (HBA) and beta (HBB) subunits from our reference assembly and aligned them to corresponding amino sequences obtained from the UniProtKB database (Fig. 4a and Supplementary Table 7). We aligned amino sequences using `Clustal Omega` v1.2.4 [105] and visualized the resulting alignment using `Mesquite` [106] (Fig. 4a).

After discovering unique substitutions in the gelada HBA, we quantified Hb-O_2_ affinity using total hemoglobin purified from hemolysates of one individual each of gelada, hamadryas baboon, Guinea baboon, anubis baboon, and human (Supplementary Table 8) using previously described methods [107]. Briefly, proteins were purified by anion-exchange FPLC, removing endogenous organic phosphates and yielding stripped samples. Using purified Hb solutions (0.4 mM heme), we measured O_2_ equilibrium curves in the absence (stripped) or presence of allosteric effectors 0.1 M KCl and 2,3-diphosphoglycerate (DPG; at 2-fold molar excess). Reactions were run at 37°C in 0.1 M HEPES buffer with 0.5 mM EDTA. *P*_50_) values were measured at three different pH levels: ~7.2, ~7.4, and ~7.7. A linear least-squares regression comparing pH and log *P*_50_) was computed and the resulting equations were used to correct *P*_50_) values to pH 7.4 for each of the gelada, baboon (three species combined), and human datasets.

We predicted that gelada hemoglobins would display increased oxygen affinity compared to those of baboons and humans. To test these predictions, we used two approaches. First, we plotted predicted *P*_50_) values at pH 7.4 for each taxon with error bars ± the standard errors of the estimates (SEEs) from the respective regression models (after exponentiation to reverse the log scales for each). We then assessed the resulting error bars for overlap, with overlap indicating no statistical difference in predicted *P*_50_) (Fig. 4b). Second, we calculated the differences in gelada vs. baboon and gelada vs. human predicted log *P*_50_) at pH 7.4 as well as the standard errors of the differences, then performed onesided *t*-tests using the equations specified by Rees & Henry [108] (case assuming homogeneity of variance; equations 3–6) to test the alternative hypotheses that gelada log *P*_50_) < baboon log *P*_50_) and gelada log *P*_50_) < human log *P*_50_). We performed these comparisons for both the stripped condition and the condition in the presence of allosteric effectors.

### Analysis of blood hemoglobin concentrations

We compared venous blood hemoglobin concentrations from this study (n=92, mean elevation ≈ 3,250 m) to corresponding measurements in zoo geladas (n=42, mean elevation ≈ 100 m) [43] and captive hamadryas baboons (n=1023, mean elevation ≈ 50 m) [44]. As the reference zoo gelada data do not differentiate sexes, we performed all comparisons with both sexes grouped together. We tested for differences of means between (1) wild geladas vs. zoo geladas and (2) wild geladas vs. captive hamadryas baboons using Welch’s *t*-test with the means, standard deviations, and sample sizes of each population as input.

For visual comparison, we plotted the means and standard deviations for hemoglobin concentration values from the Simien Mountains and zoo gelada reference values together with data from a metaanalysis of human populations across altitudes [35]. To facilitate comparison, we excluded hemoglobin concentration values from infants and juveniles and combined values between adult males and females of each human population at each reported altitude. As hemoglobin values were provided across 1 km ranges, we assigned a single elevation value as the midpoint of each 1 km range (we assigned 5,500 m for values in the category >5,000 m). We highlight differences in altitude-based hemoglobin concentrations among populations by fitting separate regression lines for (1) Tibetans and Sherpa and (2) all other human populations (Fig. 4c).

### Analysis of chest circumference

We analyzed relative chest circumference in geladas by controlling for either body mass, waist circumference, or both (Fig. 4d). We combined gelada measurement data with decades of baboon measurement data collected by two authors (J. Phillip-Conroy and C. Jolly). We restricted our analysis to adults over six years of age, estimated either from dentition [50] or calculated from known birth dates, and removed baboons of mixed species ancestry. After filtering, our comparison consisted of n=78 geladas and n=482 baboons. We tested for significantly larger lung volumes in geladas by running linear models with chest circumference as the dependent variable and sex, genus (*Papio* vs. *Theropithecus*), and an interaction term between genus and species as covariates to take into account the nested nature of species within genera. Additionally, in each of three linear models, we included (1) body mass, (2) waist circumference, or (3) both body mass and waist circumference as additional covariates to control for aspects of body size. For all models, we adjusted *P* values for one-sided hypothesis testing using the *t* distribution.

### Ortholog determination

We compared protein and gene sequences from the gelada genome to a dataset of 39 additional (40 total) mammalian genomes (Supplementary Fig. 4 and Supplementary Table 2) obtained from NCBI and Ensembl. Homologous relationships were determined using two approaches. In the first approach, we used a *de novo* orthology inference approach implemented in `OrthoFinder` [56] to assign proteins to orthogroups, which we then used to identify single-copy orthologous sequences and coding sequences under positive selection. In the second approach, we used the hidden Markov model approach implemented in `HMMER3` [109] to assign proteins to curated phylogenetically based gene families in the TreeFam9 database [57, 58], which we then used to identify expansions and contractions of gene families.

We used `OrthoFinder` to identify candidate singlecopy orthologs, using the longest translation form of each gene as inputs. We defined single-copy orthogroups as orthogroups for which the number of assigned genes in any taxon was either 0 or 1. This definition takes into account the possibility that genes are missing in some of the analyzed genomes due to incomplete assembly or annotation, resulting in 0 copies.

While the longest translation forms of each gene are useful for homolog determination as they maximize the amount of sequence information, they are not necessarily optimal for alignment as they tend to introduce nonshared exons leading to a greater number of misaligned positions. To address this problem, we used the protein alignment optimizer heuristic implemented in the software `PALO` [110] to select optimal isoforms for the analysis. Rather than selecting the longest isoform, `PALO` selects isoforms that are most similar in length. Because the `PALO` algorithm is combinatorial by nature, the computational burden increases exponentially with the number of possible length combinations to test and the software imposes an internal limit of 100 million length combinations. For orthogroups surpassing this threshold, we thus implemented a stepwise strategy in which we rank-ordered taxa according to their number of unique protein lengths, then ran `PALO` in the largest possible group of taxa for which the product of their protein counts did not exceed 100 million. After selecting one protein length per taxon for this group, we repeated the procedure as necessary until all species could be run. When multiple isoforms shared the optimum length selected by `PALO` for a given taxon, we selected one at random.

We used `HMMER3` [109] to assign proteins to gene families in the TreeFam9 database. The longest translation forms of each gene were used as inputs and the gene family with the highest bit score was assigned to each gene.

### Positive selection on protein-coding genes analysis

To identify proteins under positive selection, we generated alignments for all single-copy orthologs identified by `OrthoFinder` and using isoforms selected by `PALO`. We aligned amino acid sequences using `Clustal Omega` v1.2.4 [105], then generated codon alignments using the `pal2nal.pl` script from the `PhaME` toolkit [111]. We excluded all alignments for which (1) fewer than 36 taxa (< 90%) had sequences and (2) the total alignment was either less than 120 nucleotides (< 40 codons/amino acids) long or less than 25% the length of the full gelada protein. We then tested for positive selection using two *d*_*N*_/*d*_*S*_-based approaches: (1) branch-site models implemented in `PAML` v4.9 [54] and (2) genewide models implemented in `HyPhy` [112]. For our `PAML` analysis, we ran likelihood ratio tests on codon alignments using the “M2a” model of positive selection in the program `codeml` (`model = 2, NSsites = 2`).

For our `HyPhy` analysis, we ran likelihood ratio tests for episodic positive selection using `BUSTED` [55]. For both analyses, we used a consensus chronogram downloaded from TimeTree [24, 25] including all 40 taxa (Supplementary Fig. 4) as input into our models, with missing branches removed as necessary for each alignment. We corrected all *P* values using a Benjamini-Hochberg procedure [113] (Supplementary Table 3).

### Gene family expansion analysis

We tested for significant gene family size changes using our previously described protein assignments to the TreeFam9 [57, 58] database. Expansions and contractions were determined using `CAFE` 4.2 [59], which uses a probabilistic graphical model based on a random birth/death process to calculate the probability of transitions (*l*) in gene family size from parent to child nodes in a phylogenetic tree. For this analysis, we allowed the program to estimate the most likely value of lambda (6.09e-4) and used a consensus chronogram from TimeTree [24, 25] (Supplementary Fig. 4) as input. Because `CAFE` reports a branch-specific *P* value that indicates rapid evolution and not necessarily expansion, we defined expanded gene families as those that were both larger in *T. gelada* relative to the most recent common ancestor (MRCA) and significant at a false discovery rate (FDR) threshold of 20%. Gene families that thus exhibited significant expansion were interpreted as putative targets of selection in geladas (Supplementary Fig. 5 and Supplementary Table 5).

### Gene Ontology enrichment analyses

We performed Gene Ontology (GO) [114, 115] enrichment analyses in order to identify biological processes that are differentially associated with signatures of positive selection and gene family expansion.

For our protein positive selection analysis, we downloaded GO annotations associated with all ENSEMBL genes in our analysis, obtained using `biomaRt` [116]. For each orthogroup, we then assigned the combined, non-redundant set of GO terms and filtered to include only terms in the biological process ontology. We then tested for enrichment of low *P* values using threshold-independent Kolmogorov–Smirnov (KS) tests implemented using `topGO` [117], which corrects for the correlated nature of the GO graph network. We implemented tests in `topGO` using the “weight01” algorithm, excluding GO terms with fewer than 10 associated genes. We report enriched biological processes that passed a threshold (FDR-adjusted *P* < 0.1) using both our `PAML` and `BUSTED` *P* values analyzed separately (Supplementary Table 4).

For our gene family expansions analysis, we annotated gene families in the TreeFam9 database [58] based on provided mappings of gene families to accessioned proteins in the UniprotKB database [118]. We used the Uniprot accessions to assign proteins to ENSEMBL genes using `biomaRt` [116], then linked the combined, non-redundant set of GO terms associated with genes from the human (GRCh38) genome to each TreeFam9 family. We tested for enrichment of low *P* values using KS tests with identical settings and filters to those used in our protein positive selection analysis. We used branch-specific *P* values for the gelada branch from `CAFE` as input. Because *P* values from `CAFE` are nondirectional and ranged from 0 to 0.5, however, we first rank-ordered *P* values according to the strength of evidence for expansion by subtracting the *P* values from 1 whenever the gene family contracted in size in the gelada branch (i.e., fewer genes in the gelada branch compared to the ancestral gelada-baboon node). We considered all biological processes with an FDR-adjusted *P* < 0.1 to be significantly enriched (Supplementary Table 6).

### Gelada accelerated region analysis

We used an accelerated region approach modeled on that of Pollard *et al.* [84] on genome-wide alignment blocks to identify regions encountering accelerated evolution in the gelada lineage, which we refer to as “gelada accelerated regions”, or GARs.

We first obtained whole-genome alignment blocks for the “57 mammals EPO” dataset from ENSEMBL [119] (release 101), which includes Tgel 1.0 and 56 additional mammalian genomes in multiple alignment format (MAF). We subsetted alignment blocks to include only terminal branches (i.e., excluding ancestral sequences), then preprocessed MAF files using `mafTools` [120] to remove duplicate species (`mafDuplicateFilter`), set human (*Homo sapiens*) as the reference species (`mafRowOrderer`), index all blocks to the positive strand on the reference (`mafStrander`), and to sort blocks by position (`mafSorter`). We subsetted blocks to include only gelada and the species trio of human/mouse/rat, then performed local realignment within MAF blocks using `MAFFT` [121, 122] to correct misalignments. We next used `MafFilter` [123, 124] to define and extract conserved alignment blocks that met the following criteria within the human/mouse/rat species trio: (1) a block length of ≥ 50 bp, (2) gaps in no more than 10% of positions within a 50 bp window, and (3) variable sites (including gaps) in no more than 10% of positions within a 50 bp window. We used these criteria because they were within the range of effective parameters evaluated by Pollard *et al.* [84], but maximized the number of genomic regions available for downstream analyses. We retained blocks encompassing the most inclusive set of coordinate positions that passed these criteria.

We filtered all alignment blocks by the criteria described above using `AlnFilter` and `EntropyFilter` from the `MafFilter` software package [124]. Notably, both of these algorithms are designed to identify and remove sites failing filters across sliding windows. Blocks are then normally split to remove sites failing filters within any window and trimmed blocks containing residual coordinates are returned as output. Because our pipeline instead required identifying and retaining sites passing filters, we modified and recompiled the source code of `MafFilter` and its `Bio++` dependencies [125, 126] to direct windows failing filters to the output and windows passing filters to the “trash”. In so doing, we took advantage of a feature of `MafFilter` by which windows failing filters are optionally redirected to a “trash” MAF file, with adjacent coordinate sites merged into contiguous blocks for perusal. By directing coordinates passing filters to the “trash” and by using the resulting blocks as inputs for the remainder of the pipeline, we were able to induce the desired behavior from the software.

After identifying sites passing filters, we extracted their coordinates using `MafFilter OutputCoordinates` and used the resulting file to extract the corresponding positions from the MAF blocks containing all species using maf_parse from the `PHAST` software package [127]. We then repeated local alignment of MAF blocks using `MAFFT`and indexed blocks to the gelada reference genome using `mafRowOrderer` and `mafStrander` from `mafTools` to create the final MAF blocks. A total of 60,345 blocks passing filters were included in our analysis.

We tested for acceleration within blocks using the CONS model described by Pollard *et al.* [84] and the phylogenetic tree included with the ENSEMBL “59 mammals EPO” dataset. The CONS model fits a general time-reversible model (REV) on aligned sequences using `phyloFit` from the `PHAST` v1.5 software package [127]. Acceleration is assessed by a likelihood ratio test (LRT), comparing a phylogenetic model in which branches are scaled across the tree in equal proportions and a model in which the foreground branch (gelada) is scaled separately from the remainder of the tree. The LRT statistic is the log ratio of the likelihood of the latter (alternate) model to the former (null) model multiplied by 2. We calculated significance from the LRT statistic using the chi-squared distribution.

To assess the distribution of acceleration scores within blocks, we also ran `phyloP` from the `PHAST` v1.5 package, which tests for acceleration or conservation at the nucleotide level. We ran `phyloP` for all 60,345 blocks in our analysis with the same phylogenetic model (REV) and phylogenetic tree, with the *P*-value reporting mode set to “CONACC” to distinguish between signals of conservation and acceleration. We then extracted genomic positions and CONACC *P* values from the `phyloP` output files.

We defined GARs as blocks for which FDR-adjusted *P* < 0.2 from the CONS model. A total of 29 blocks passed this threshold (Supplementary Fig. 6 and Supplementary Table 9), which we classified as either exonic, intronic, or intergenic based on overlap with annotated regions in the ENSEMBL GFF3 file. To identify biological processes associated with these blocks, we matched all 60,345 blocks passing filters to their nearest genes using `GenomicRanges` [128] in `R`, then downloaded all associated GO biological processes to genes using `biomaRt` [116]. To identify candidate regulatory elements, we also matched blocks with overlapping ChromHMM chromatin state annotations (15-state model, 127 epigenomes) obtained from the Roadmap Consortium [129]. We focused on 8 states that show putatively regulatory hallmarks (i.e., enrichment of ChIP-seq binding sites and enrichment of DNase peaks): active transcription start site (TSS), flanking active TSS, transcribed at genes 5’ and 3’, genic enhancers, enhancers, bivalent/poised TSS, flanking bivalent/poised TSS, and bivalent enhancer.

For two pairs of GARs (GAR26–GAR27 and GAR28–GAR29) that were nearest to the same genes, we estimated linkage disequilibrium between each GAR within each pair. To perform this analysis, we used whole-genome gelada variant data (described below) and calculated *r*^*2*^ between sites using `VCFtools` v0.1.16 [130]. We limited our sample to geladas, excluded indels and non-biallelic SNVs, and filtered to only sites within the boundaries of each GAR ± 1000 bp. We calculated *r*^*2*^ between all pairs of sites between GARs by setting a minimum distance of 500 kb between sites (arguments: `-geno-r2-ld-window-bp-min 500000`), then calculated mean *r*^*2*^ across all pairs of sites.

### Whole-genome population resequencing and analysis

#### Library generation and sequencing

DNA was extracted from whole-blood samples or muscle samples using the DNeasy Blood & Tissue Kit (QIAGEN), following manufacturer recommendations for maximizing yield and quality. Concentration was assessed by Qubit 3 (Invitrogen) and 50 ng of DNA were used as input for whole-genome sequencing (WGS). Libraries were prepared using the Nextera DNA Library Prep protocol (Illumina). Briefly, DNA was added to a 10 μl reaction containing 5 μl of TD buffer and 1 μl of tagment DNA enzyme (TDE1), then incubated at 55°C for 5 minutes. Tagmentation reactions were cleaned using 2x concentration of Ampure XP beads (Beckman Coulter), then 10 μl of cleaned DNA were added to a 24 μl PCR reaction including 1x NEBNext Q5 master mix (New England Biolabs) and 0.42 μM each of indexed P5/P7 primers. Libraries were amplified using six cycles of PCR and cleaned using 0.65x concentration of Ampure XP beads (Beckman Coulter). Libraries were pooled equimolarly and sequenced on either the Illumina HiSeq X or NovaSeq 6000 platforms (2×151 bp sequencing) to a median coverage of 11.54x.

#### Mapping and genotyping

We mapped reads to either the gelada reference genome (Tgel 1.0) or the anubis baboon reference genome (Panubis 1.0) [131] using the `speedseq align` v0.1.2 pipeline [132], which includes reference mapping with `BWA-MEM` [133], duplicate marking and discordant-read/split-read extraction with `SAMBLASTER` [134], and position sorting and BAM file indexing with `SAMBAMBA` [135].

We genotyped reads using a pipeline implemented in `GATK` v4.1.2.0. We genotyped on a per-sample basis using `GATK HaplotypeCaller` to generate GVCF files. We then performed joint genotyping across samples using `GATK GenotypeGVCFs`, after first creating a GenomicsDB workspace using `GATK GenomicsDBImport`. We filtered variants using `GATK VariantFiltration` with the filters “`QD < 2.0, MQ < 40.0, FS > 60.0, MQRankSum < −12.5, ReadPosRankSum < −8.0`, and `SOR > 3.0`”, then extracted sites passing filters using `BCFtools` [136, 137].

We used the resulting genotypes to recalibrate base quality scores using the `GATK BaseRecalibrator` and `ApplyBQSR` workflows. We then repeated persample variant calling, joint genotyping, and variant filtration to sequentially improve our genotype qualities. We performed a total of two rounds of base quality score recalibration bootstrapping in this manner, then repeated our genotyping pipeline a final time to generate final genotypes in VCF format. Our final VCFs included 48,744,921 variants mapped to Tgel 1.0 (chromosomes 1–21 and X) and 35,615,864 variants mapped to Panubis 1.0 (chromosomes 1–20, X, and Y).

#### Population structure analysis

Separately from our `GATK` genotyping pipeline, we implemented the genotyping uncertainty models in `ANGSD` [138] and `PCAngsd` [34] to analyze population structure. We used BAM files mapped to the gelada genome (Tgel 1.0) as input. We then used `ANGSD` v0.921 to generate genotype likelihoods in beagle format (arguments: `-GL 1 -doGlf 2 -doMajorMinor 1 -doMaf 2 -minMaf 0.05 -SNP_pval 1e-6 -minQ 20 -minMapQ 30 -skipTriallelic 1 -minInd 22 -doDepth 1 -doCounts 1`). We then used `PCAngsd` v0.95 to estimate admixture proportions (arguments: `-admix -admix_alpha 5000`).

#### Determination of geographic provenience

To determine the provenience of the 17 zoo individuals in our study, we assembled mitochondrial DNA sequences from WGS reads for all individuals in our dataset and aligned them to the mitochondrial DNA dataset of Zinner *et al.* [6], which consists of cytochrome *b* + hypervariable region I (HVI) D-loop sequences from wild geladas across their natural distribution. We assembled complete mitochondrial sequences using `GetOrganelle` v1.7.5 [139], which uses `Bowtie2` [140], `BLAST` [141], and `SPAdes` [142] to assemble circular genomes *de novo* from WGS data. To reduce computational burden, we limited our input to 2 million read pairs (~600 Mb) per individual randomly sampled using `seqtk` v1.3 [143] and incorporated a reference mitochondrial genome assembly (GenBank accession FJ785426.1) [144] as an input seed sequence. We then extracted the cytochrome *b* + HVI region for each sample by aligning to Zinner *et al.* [6] sequences using `EMBOSS water` v6.6.0 [145].

We combined all new gelada cytochrome *b* + HVI sequences (Supplementary Table 1) with 61 gelada haplotypes from Zinner *et al.* [6], one sequence from a hamadryas baboon in our dataset (FIL001), and one sequence from a rhesus macaque (GenBank accession NC_005943.1) [146]. We then aligned all nonredundant haplotypes using `Clustal Omega` v1.2.4 [105]. To infer a phylogenetic tree, we ran `IQ-TREE` v2.1.2 [147] with two partitions, protein-coding (1–1134) and non-coding (1135–1737) sequences, and set the rhesus macaque sequence as the outgroup. We used the `ModelFinder` option [148] within `IQ-TREE` to select the best nucleotide substitution models according to the the Bayesian Information Criterion (HKY+F+G4 and HKY+F+I+G4 for protein-coding and non-coding partitions, respectively) and ran 10,000 ultrafast bootstrap replicates (Supplementary Fig. 7).

#### Demographic history

We estimated demographic histories using `MSMC2` v.2.1.2 [149, 150] (Fig. 3a). We used per-sample VCFs, described earlier, and generated a mask file to exclude sites with excessively low or high coverage. We used `Mosdepth` [151] to calculate mean sample coverage and to generate a BED file per-sample marking sites with sequencing depth above a minimum of 50% mean coverage and below a maximum of 250% mean coverage. We then merged VCF files and mask files using the `generate_multihetsep.py` script and ran `MSMC2` using the resulting file as input. We set 11.67 years as the average generation time, which we derived by calculating the average maternal age at birth from the SMGRP longitudinal life history database, and 0.5×10^−8^ as the mutation rate (*μ*), which is derived from estimates in anubis baboons [152]. For plotting, we excluded samples with < 10x mean coverage.

### Analysis of heterozygosity and runs of homozygosity

We calculated average heterozygosity and identified runs of homozygosity for 90 individuals from the Central and Northern gelada populations, a captive gelada group, and a population of hamadryas baboons (*Papio hamadryas*) from Filoha, Ethiopia (Fig. 3b). We used variants called from data mapped to the anubis baboon reference genome (Panubis 1.0) [131]. Heterozygosity was calculated as per-site average in 100 kb windows with a 10 kb slide across all autosomes in the Panubis 1.0 reference for each individual. Windows were generated with `BEDtools makewindows` v2.29.2 [153] and number and percent callable sites within each window were identified with `BEDtools intersect` v1.10.2 [153]. A window was considered part of a run of homozygosity if its average heterozygosity was below 0.0002. We identified runs of adjacent windows with Ho < 0.0002 with the `rle` function in `R` v3.6.0 [154] and calculated the number of callable bases contained within runs of homozygosity <1 Mb, 1–3 Mb, and >3 Mb in length.

