## Supplementary Figures for "High-altitude adaptation and incipient speciation in geladas"

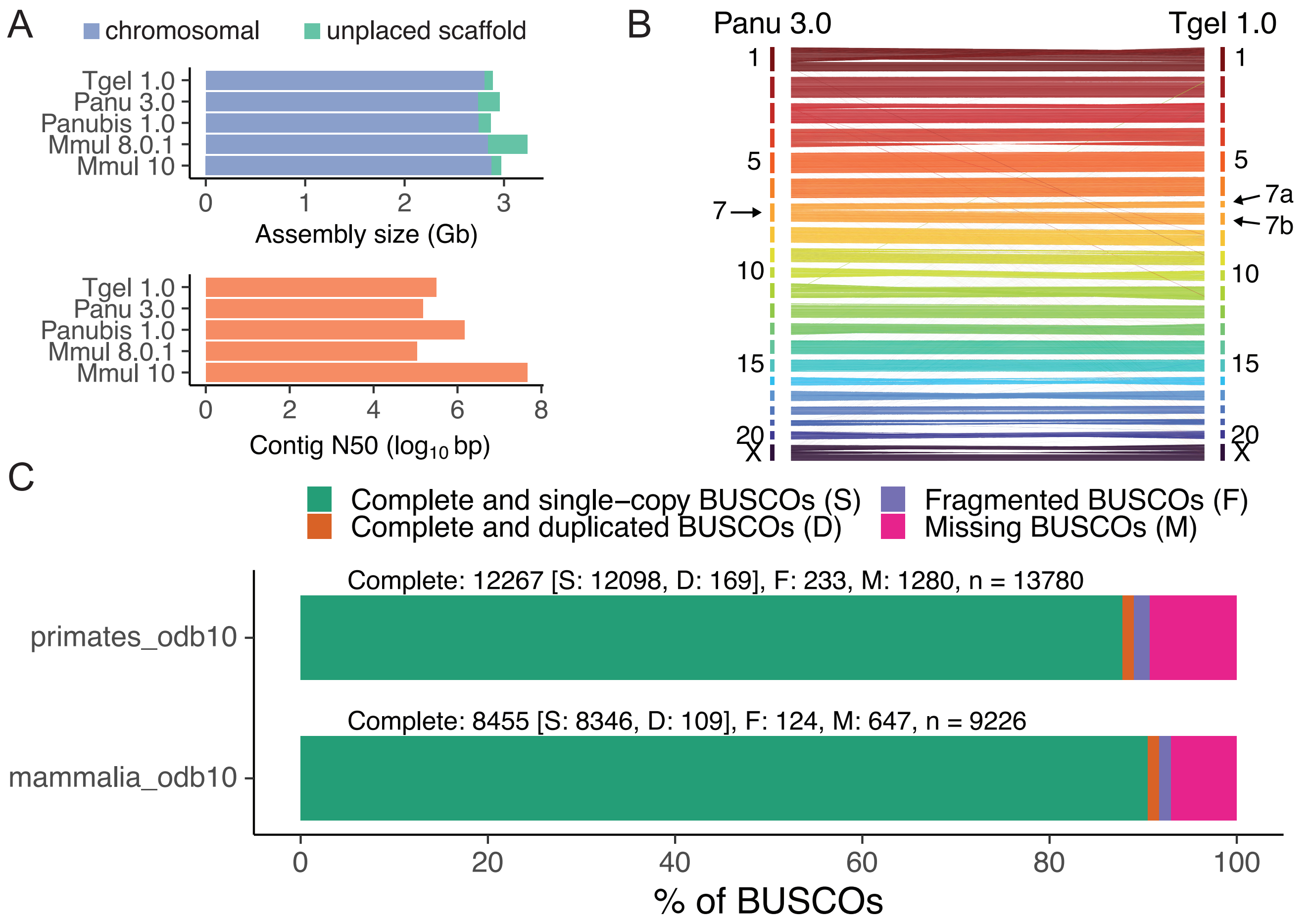

**Supplementary Figure 1. Gelada reference assembly quality and synteny.** (A) The gelada reference assembly (Tgel 1.0) compared to closely related papionin assemblies in assembly size (top) and contig N50 (bottom). (B) Synteny links between the anubis baboon (Panu 3.0) and gelada (Tgel 1.0) assemblies reveal strong collinearity between genomes. (C) BUSCO analysis of the gelada reference assembly reveals a relatively intact and complete assembly.

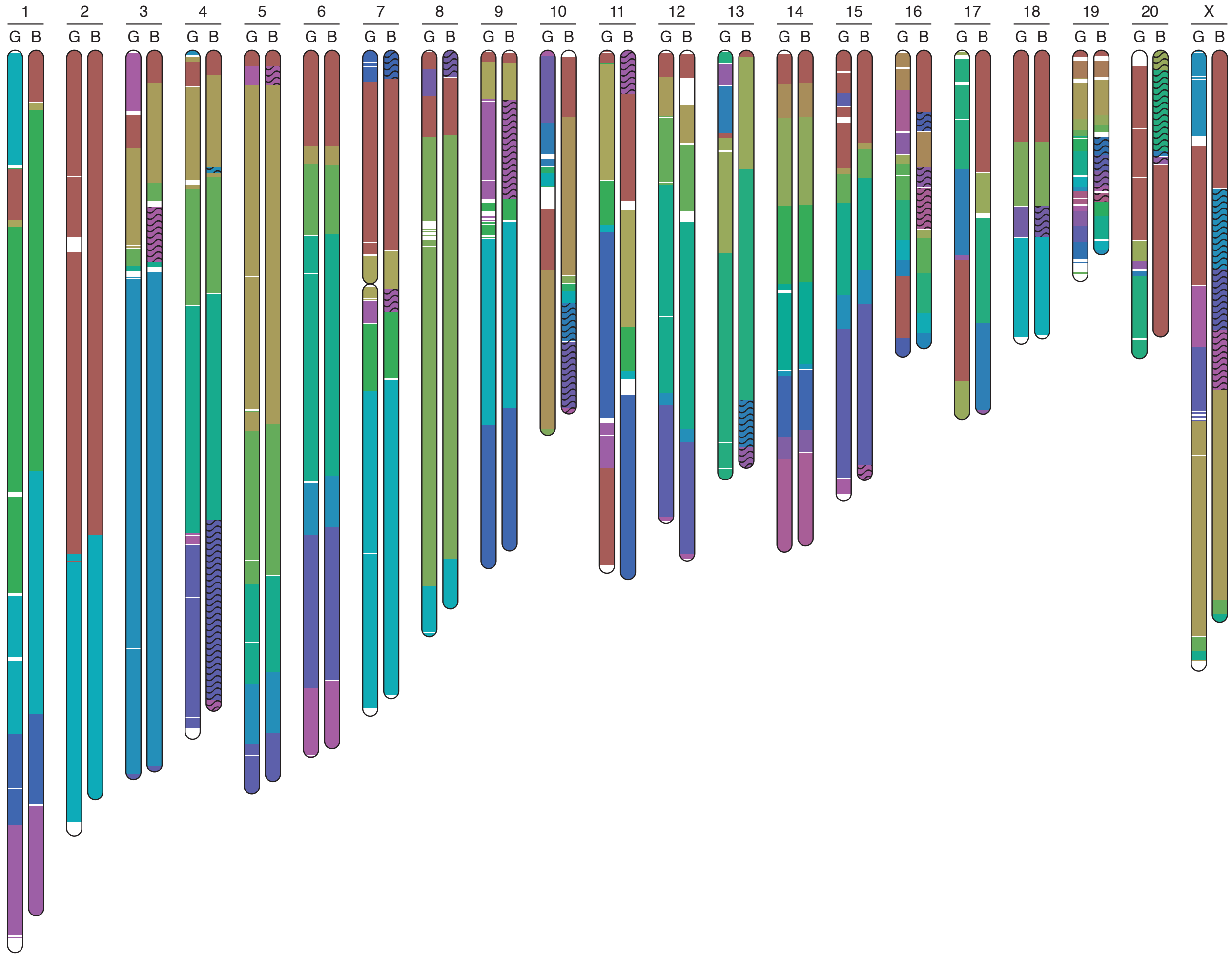

**Supplementary Figure 2. Synteny blocks and chromosomal rearrangements in the gelada reference assembly.** Ideogram generated using the alignment-free method implemented in SMASH reveals synteny blocks as well as chromosomal rearrangements between gelada ("G", Tgel 1.0) and anubis baboon ("B", Panu 3.0) genomes.

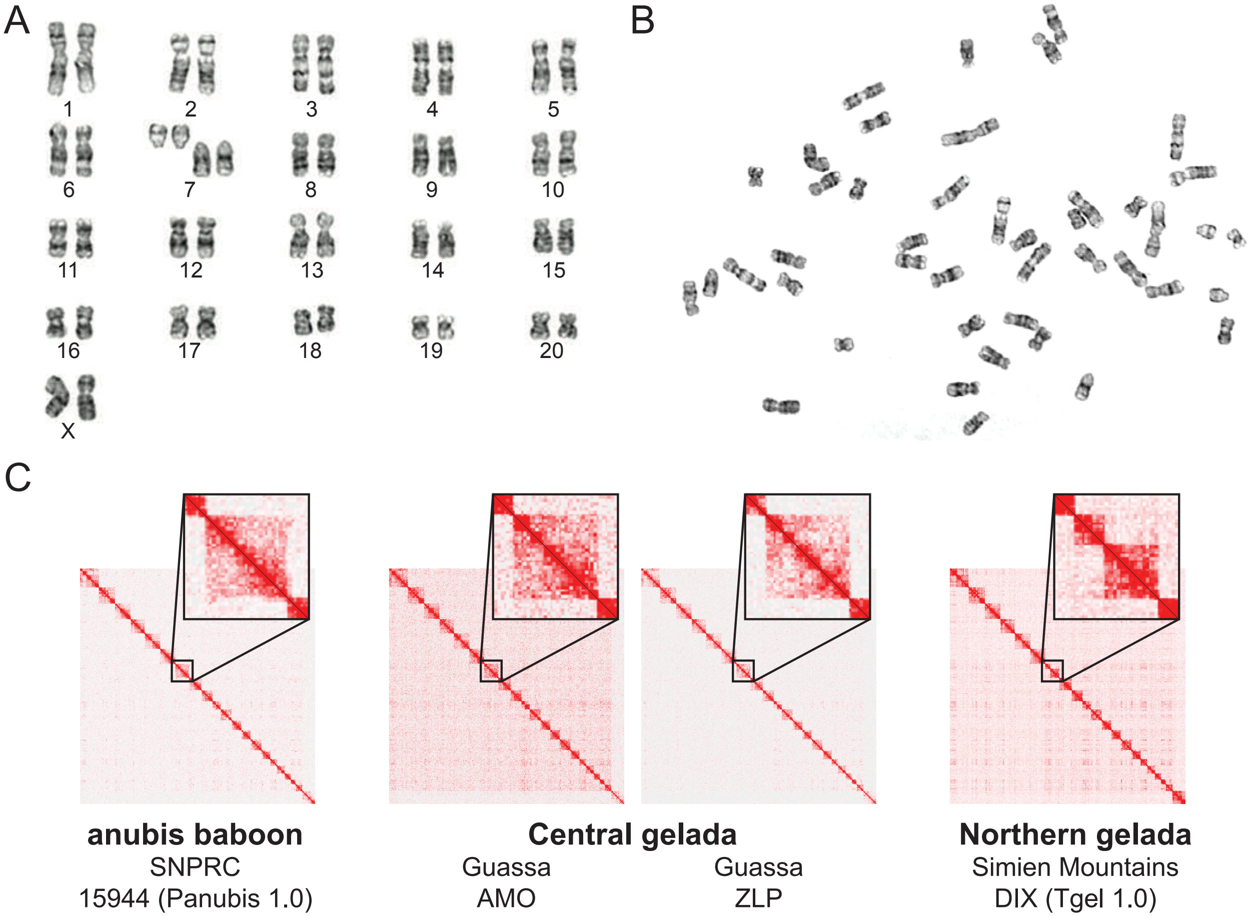

**Supplementary Figure 3. Karyotyping and a unique centric fission in gelada chromosome 7.** (A) Full karyotype of our female reference individual (DIX). (B) Example G-banded chromosome spread with 44 counted chromosomes. (C) Analysis of Hi-C libraries allows for determination of the presence/absence of a centric fission in chromosome 7 without the need for live cells, which are difficult to obtain from wild populations. Two wild Central gelada individuals showed abundant contacts between the two arms of chromosome 7, indicating an intact chromosome and providing the first provenienced sampling to our knowledge of Central gelada karyotypes.

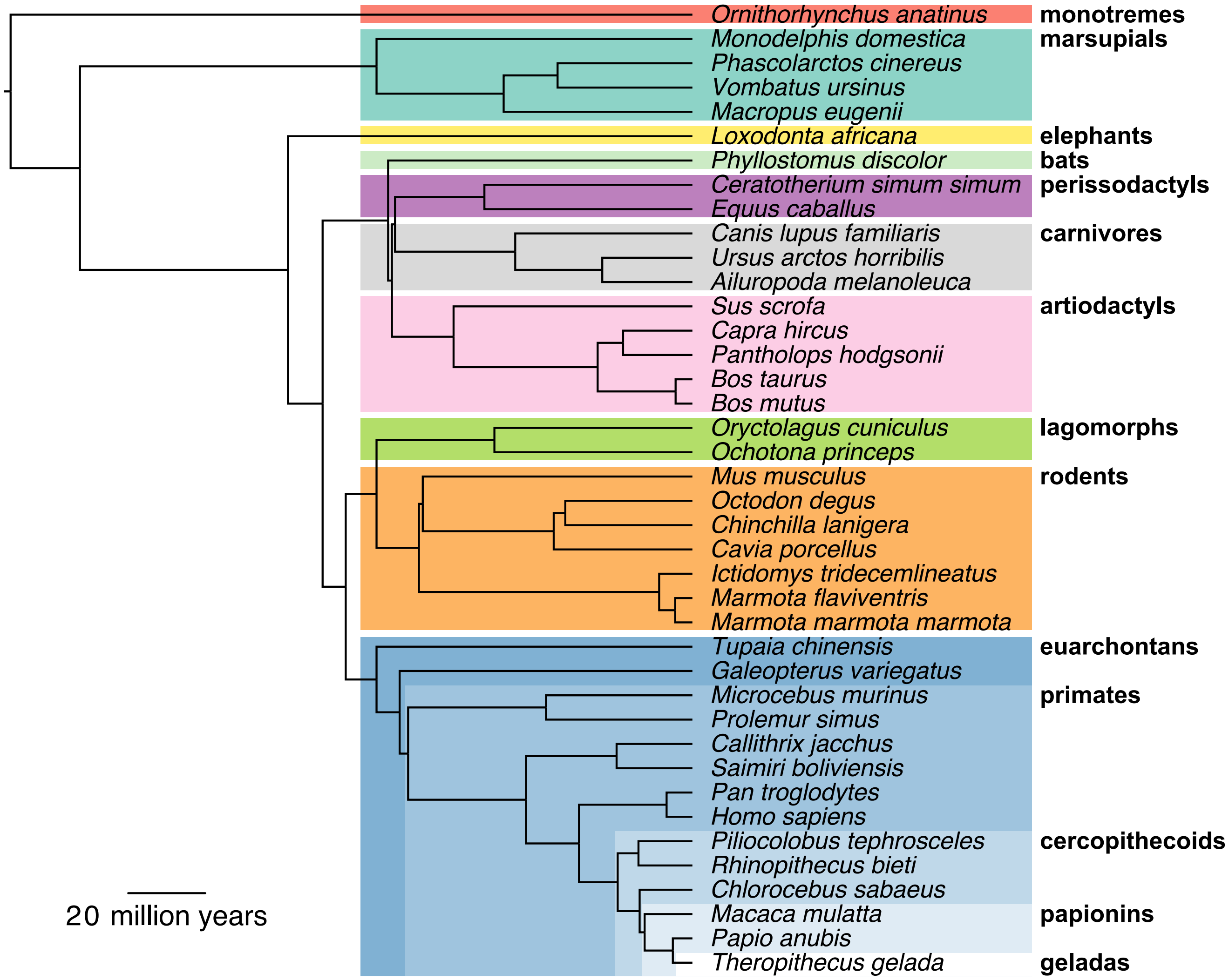

**Supplementary Figure 4. Genome assemblies included in positive selection and gene family expansions analyses.** Chronogram was obtained from TimeTree [24,25].

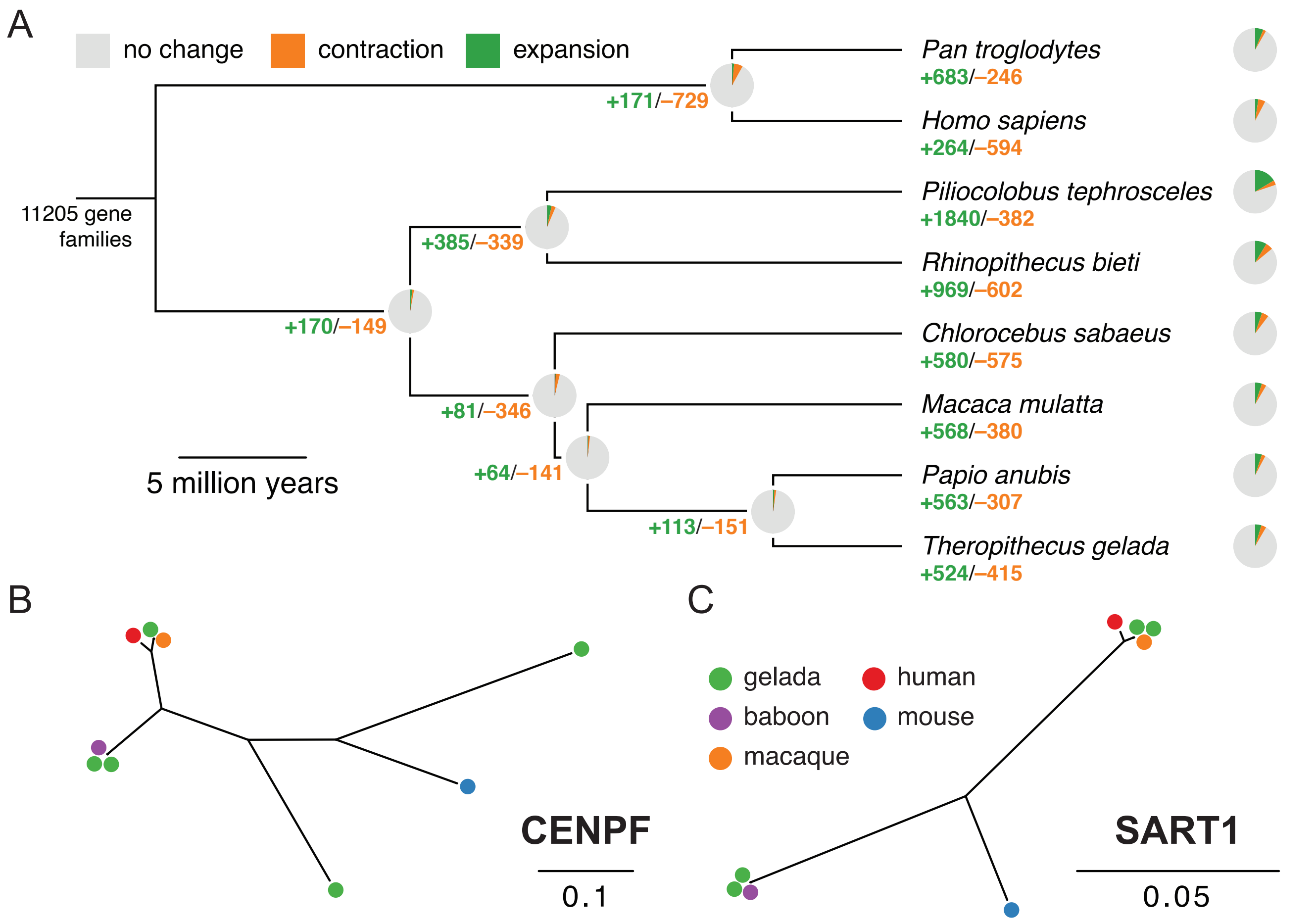

**Supplementary Figure 5. Gene family size changes in the gelada genome.** (A) Gene family expansions and contractions across the catarrhine tree. Here, expansion and contraction estimates are from CAFE and do not use the more stringent statistical thresholds used for downstream analyses. (B) Example of a significantly expanded gene family containing *CENPF*, which is found with five copies in geladas. (C) Example of a significantly expanded gene family containing *SART1*, which is found with four copies in geladas. Proteins are grouped using a neighbor-joining tree.

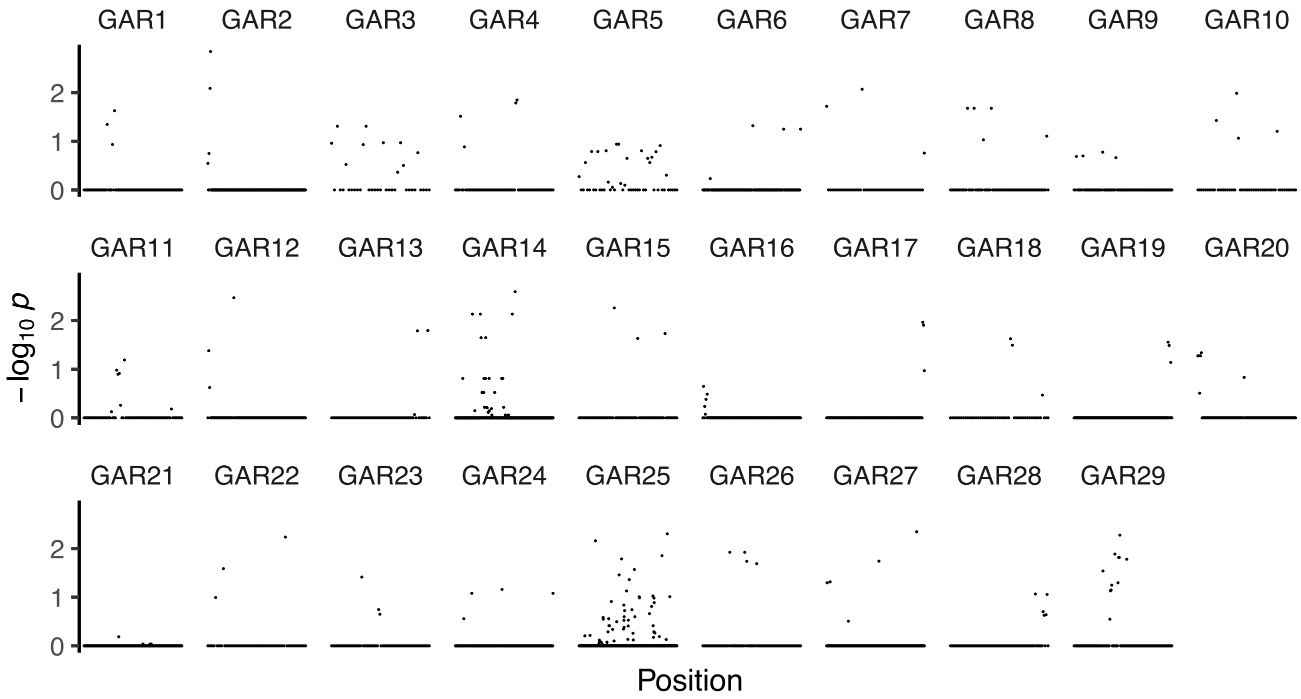

**Supplementary Figure 6. Robust signals of acceleration across GARs.** Per-base acceleration scores estimated with phyloP reveal the distribution of elevated signals of acceleration across GARs. Some GARs (e.g., GAR16 and GAR 17) show signals of acceleration that are highly localized while other GARs (e.g., GAR5, GAR14, and GAR25) show numerous changes that are more uniform across larger regions.

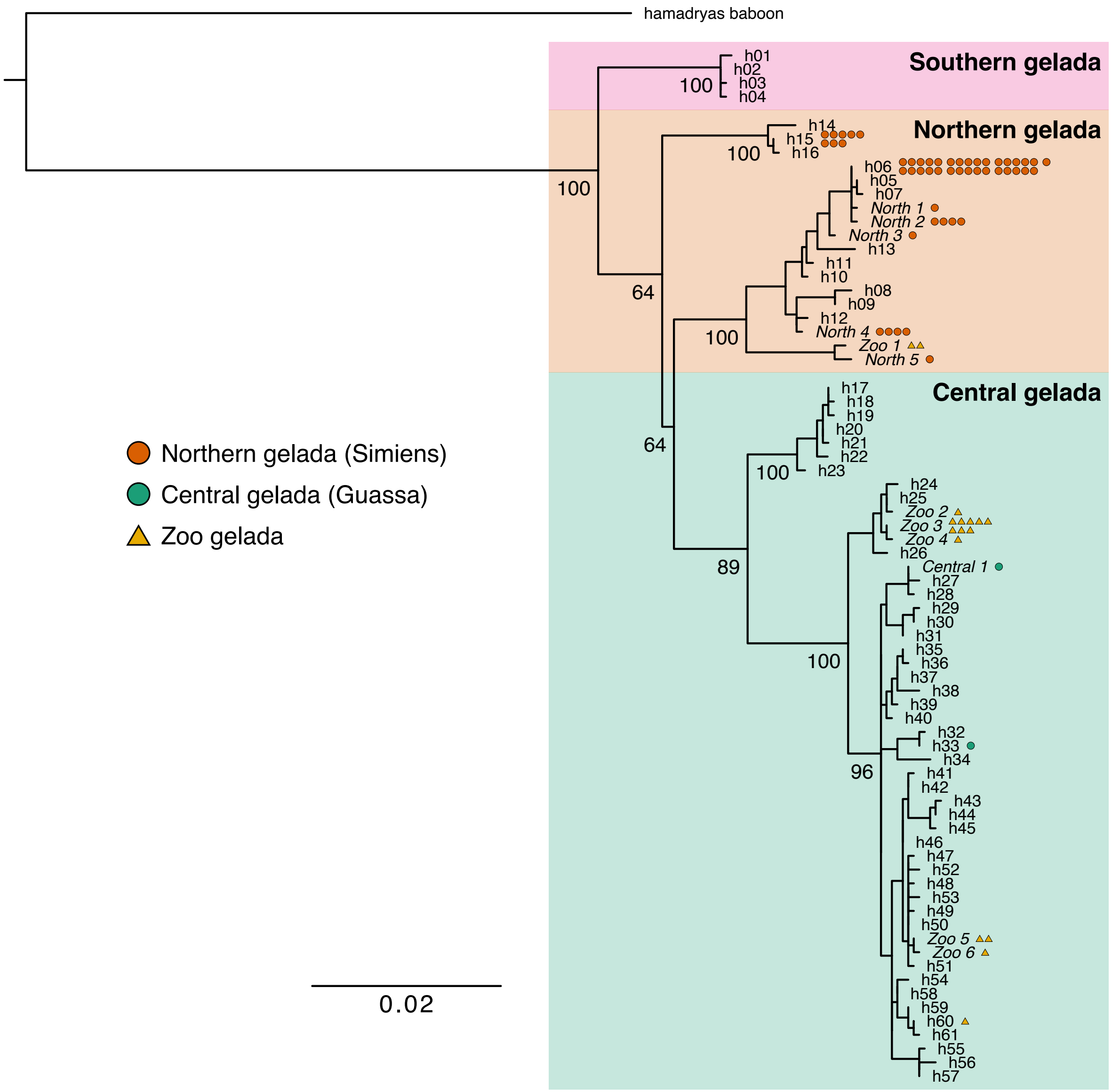

**Supplementary Figure 7. Provenience of captive gelada samples.** Maximum likelihood phylogenetic tree from the cytochrome *b* + hypervariable region I (HVI) D-loop mitochondrial region informs on the geographic origin of geladas in zoos. Individuals in our study were assigned either to gelada haplotypes determined by Zinner *et al.* [6] (h01–h61) or to new haplotypes determined in the current study (labeled in italics; Supplementary Table 1). Gelada individuals sampled from the wild were exclusively assigned to clades matching their geographic origin (Northern or Central). Zoo individuals were assigned to the Central clade with the exception of a single haplotype shared by two zoo individuals, which was assigned to the Northern clade. The two zoo individuals both have heterozygous (2n=43) karyotypes and elevated fractions of Northern genome-wide ancestry, indicating that they likely descended from a Northern individual. A rhesus macaque reference sequence (GenBank accession NC_005943.1) was used to root the tree and is not shown. Bootstrap support values are shown for major nodes.
